# A longitudinal CA2/3 to CA1 circuit for initiating context-dependent associative learning

**DOI:** 10.1101/2022.02.05.479215

**Authors:** Hao-Shan Chen, Shou Qiu, Guang-Ling Wang, Rong-Rong Yang, Na Zhang, Jin-Ni Wu, Qi-Xin Yang, Chun Xu

## Abstract

Associating environmental context with emotional experience is a vital brain function that requires the activity in both dorsal and ventral hippocampus. While only ventral hippocampus connects with the amygdala, a hub for fear learning, it remains unclear how the two hippocampal areas interact during contextual fear conditioning (CFC). We found that projections from dorsal CA2/CA3 (dCA2/3) to the dorsal part of ventral CA1 (vCA1d) contributed significantly to CFC. Deep-brain Ca^2+^ imaging revealed a CFC-enhanced difference in neuronal activities evoked by conditioned vs. neutral environmental context in both dCA2/3 and vCA1d areas. Notably, contextual fear retrieval correlated with changes in conditioned context-evoked activity in vCA1d, but not in dCA2/3. Furthermore, slice recordings showed that CFC potentiated dCA2/3-to-vCA1d projection strength, in line with coordinated elevation of monosynaptic excitation and reduction of disynaptic inhibition mediated by somatostatin-expressing interneurons. Thus, vCA1d represents a critical site for initiating emotional association with contextual information from dCA2/3.

## Introduction

Associative learning enables adaptive behaviors of the animal in different environmental contexts in a manner that depends on context-associated emotional experiences (Fanselow and Poulos, 2005; LeDoux, 2000; Maren, 2001; Maren et al., 2013; Tovote et al., 2015). The hippocampus is known to be critical for context-dependent memory formation (Cai et al., 2016; Fanselow, 2000; Goshen et al., 2011; Liu et al., 2012; Lovett-Barron et al., 2014; Reijmers et al., 2007; Zhou et al., 2017), and plays pivotal roles in episodic memory (Scoville and Milner, 1957) and spatial navigation (O’Keefe and Nadel, 1978). The rodent hippocampus is an elongated C-shaped structure extending from the septal (dorsal) to the temporal (ventral) pole. While the trisynaptic and unidirectional projections of the dentate gyrus (DG)→CA3→CA1 core circuit is present throughout the dorsal-ventral axis of the hippocampus (Andersen et al., 1971), there is mounting evidence indicating the anatomical and functional distinction between dorsal and ventral hippocampus (Fanselow and Dong, 2010). Anatomically, dorsal hippocampus mainly connects with cortical areas, whereas ventral hippocampus is more closely connected to subcortical structures such as amygdala and hypothalamus (Pitkanen et al., 2000; Swanson and Cowan, 1977). Functionally, dorsal and ventral hippocampal areas are preferentially involved in cognitive and emotional processing, respectively (Strange et al., 2014). For instance, place cells for spatial processing are more abundant and have a higher spatial selectivity in the dorsal hippocampus than those in the ventral hippocampus (Jung et al., 1994; Kjelstrup et al., 2008).

The dorsal and ventral areas of the hippocampus are both necessary for contextual fear conditioning (CFC) and context processing (Bast et al., 2001; Kim and Fanselow, 1992; Phillips and LeDoux, 1992). More recent studies showed that place cells in the dorsal CA1 and CA3 (dCA1 and dCA3) process context information by context-dependent remapping of their place fields (Hainmueller and Bartos, 2018; Leutgeb et al., 2007). However, the dorsal hippocampus does not connect directly with the amygdala, the major hub for associative fear learning (Tovote et al., 2015). This raises the question of how dorsal hippocampus transmits the context information to the amygdala in context-dependent fear learning. The ventral hippocampus and the amygdala are strongly connected (Pitkanen et al., 2000) and the projection from ventral CA1 (vCA1) to amygdala is essential for contextual fear learning and retrieval (Kheirbek et al., 2013; Kim and Cho, 2020; Xu et al., 2016). The vCA1 is thus in an ideal position to bridge the information flow from the dorsal hippocampus to the amygdala. In line with this idea, the CA3, an upstream area of CA1, extends diffuse Shaffer collaterals along the longitudinal axis (Amaral and Witter, 1989; Swanson et al., 1978). Thus, the dCA3-vCA1 projection could play a critical role in mediating the information flow from dorsal to ventral hippocampus during contextual fear learning.

Using a combination of circuit tracing, optogenetics, slice electrophysiology, photometry, single-unit recording, and deep-brain Ca^2+^ imaging, we have identified a specific pathway from the dorsal CA2/CA3 (dCA2/3) to the dorsal area of ventral CA1 (vCA1d) that is critical for CFC. We showed that CFC enhanced the difference of neuronal activities between conditioned and neutral contexts in both dCA2/3 and vCA1d areas. However, we found that contextual fear retrieval correlated with the changes of conditioned context-evoked neuronal activity only in vCA1d, but not in dCA2/3. In this dCA2/3→vCA1d circuit, we found that pyramidal cells in vCA1d received both monosynaptic glutamatergic dCA2/3 projection and feed-forward inhibition driven by the dCA2/3 projection to vCA1d interneurons. We further showed that the strength of this dCA2/3→vCA1d projection was significantly potentiated after CFC, and this potentiation involved coordinated strengthening of the monosynaptic connection and weakening of feed-forward inhibition mediated by somatostatin (SOM)-expressing interneurons. Taken together, our findings demonstrate that longitudinal dCA2/3→vCA1d pathway is essential for contextual information flow, and vCA1d is a critical site for initiating the association between context information and emotional experience.

## Results

### Longitudinal projection from dCA2/3 to vCA1

To characterize the longitudinal projection from dCA3 to vCA1, we performed retrograde tracing from vCA1 by injecting red retrobeads. To cover the large area of vCA1, we locally injected retrobeads into dorsal, median and ventral parts of vCA1 (vCA1d, vCA1m and vCA1v), respectively. These retrograde tracings yielded comparable numbers of presynaptic cells in the hippocampus. Interestingly, we found that dCA3 cells were preferentially labeled by retrograde tracing from vCA1d, but not vCA1m or vCA1v (Fig. 1a-d; Fig. S1). Notably, the retrobeads-labeled cells were found mostly in the distal part of dCA3 adjacent to CA2 area. To clarify the identity of the presynaptic cells to vCA1d, we performed immunostaining of CA2 molecular maker PCP4 (Purkinje cell protein 4) in the brain slices, in combination with retrograde tracing from vCA1d (Fig. S2a-c). The co- localization analysis revealed that nearly 1/3 vCA1d-projecting cells were positive for CA2 markers PCP4 (Fig. S2d). Therefore, we hereafter refer all vCA1d-projecting cells in the dorsal hippocampus as dCA2/3 projection cells.

**Figure 1.**
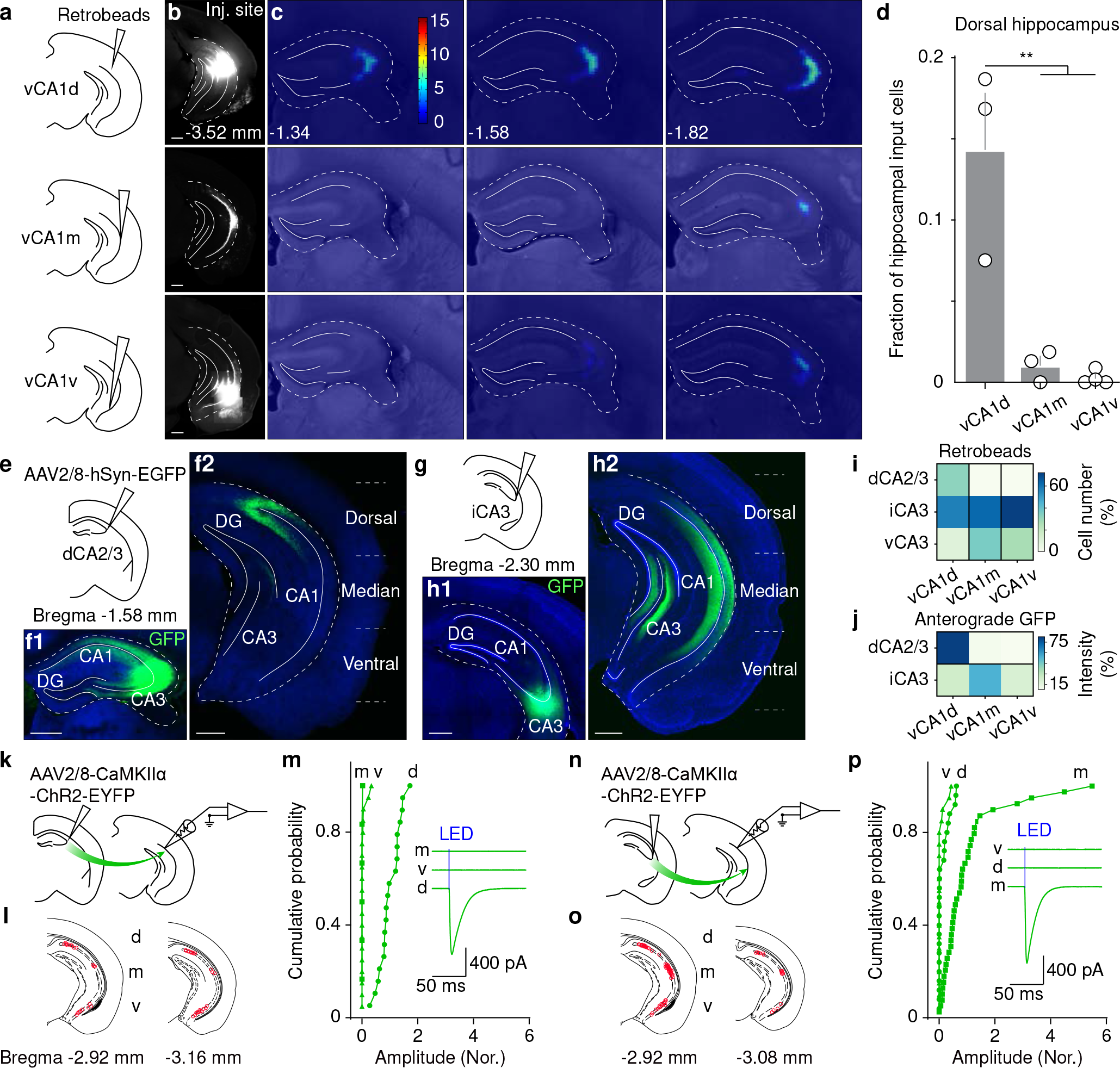
The dCA2/3 specifically targets dorsal area of vCA1 in the longitudinal axis. (a) Scheme illustrating injection of red retrobeads into vCA1d, vCA1m and vCA1v, respectively. (b and c) Example showing local injection sites (b) and density maps (arbitrary units) of retrobeads labeling in the dorsal hippocampus (c). (d) Summary (mean ± SEM) of proportion of retrobeads labeled cells in the dorsal hippocampus after injections into vCA1d (0.14 ± 0.03, N = 3 mice), vCA1m (0.01 ± 0.01, N = 3 mice) and vCA1v (0.003 ± 0.002, N = 4 mice), respectively. One-way ANOVA, F(2,7) = 18.84, P = 0.0015; Turkey’s multiple comparisons test, vCA1d vs. vCA1m, P = 0.0038; vCA1d vs. vCA1v, P = 0.0019). (e and g) Scheme illustrating injections of AAV vectors into dCA2/3 (e) and iCA3 (g), respectively. (f and h) Example showing GFP fluorescence at injection sites and downstream areas in vCA1. (i) Fractions (%) of presynaptic cell numbers labeled by retrobeads from vCA1 subregions. From vCA1d: dCA2/3, 32.7 ± 12.6%; iCA3, 57.0 ± 7.7%; vCA3, 10.3 ± 5.3%; N = 3 mice; One-way ANOVA, F(2,6) = 6.66, P = 0.03; Turkey’s multiple comparisons test, dCA2/3 vs. iCA3, P = 0.22; dCA2/3 vs. vCA3, P = 0.26. From vCA1m: dCA2/3, 0.88 ± 0.57%; iCA3, 68.8 ± 4.8%; vCA3, 30.3 ± 4.7%; N = 3 mice; One-way ANOVA, F(2,6) = 75.63, P = 5.6×10^-5^; Turkey’s multiple comparisons test, dCA2/3 vs. iCA3, P = 5.4×10^-5^; dCA2/3 vs. vCA3, P = 0.0018. From vCA1v: dCA2/3, 0.78 ± 0.73%; iCA3, 71.9 ± 7.0%; vCA3, 27.3 ± 6.9%; N = 4 mice; One-way ANOVA, F(2,9) = 39.7, P = 3.4×10^-5^; Turkey’s multiple comparisons test, dCA2/3 vs. iCA3, P = 2.7×10^-5^; dCA2/3 vs. vCA3, P = 0.023. (j) Fractions (%) of GFP intensity in the vCA1 subregions of axons from dCA2/3 and iCA3. From dCA2/3: vCA1d, 81.9 ± 4.1%; vCA1m, 9.9 ± 1.0%; vCA1v, 8.3 ± 3.2%; N = 4 mice; One-way ANOVA, F(2,9) = 188.5, P = 4.5×10^-8^; Turkey’s multiple comparisons test, vCA1d vs. vCA1m, P = 1.3×10^-7^; vCA1d vs. vCA1v, P = 1.1×10^-7^. From iCA3: vCA1d, 24.7 ± 2.0%; vCA1m, 54.4 ± 3.1%; vCA1v, 20.9 ± 2.4%; N = 4 mice; One-way ANOVA, F(2,9) = 51.33, P = 1.2×10^-5^; Turkey’s multiple comparisons test, vCA1m vs. vCA1d, P = 4.8×10^-5^; vCA1m vs. vCA1v, P = 1.8×10^-5^. (k and n) Scheme illustrating injections of AAV vectors into dCA2/3 (k) and iCA3 (n) respectively and recordings in vCA1. (l and o) Locations of recorded cells (red circles) in acute brain slices of vCA1. (m) The cumulative distribution of EPSC amplitudes (normalized by the mean amplitudes of cells in vCA1d) recorded in vCA1 cells evoked by optogenetic stimulation of dCA2/3 axons (vCA1d, 1.0 ± 0.1, n = 19 cells, N = 6 mice; vCA1m, 0.004 ± 0.004, n = 6 cells, N = 2 mice; vCA1v, 0.036 ± 0.02, n = 20 cells, N = 6 mice). Inset shows the example traces recorded from cells in vCA1d (d), vCA1m (m) and vCA1v (v), respectively. (p) The cumulative distribution of EPSC amplitudes (normalized by the mean amplitudes of cells in vCA1m) recorded in vCA1 cells evoked by optogenetic stimulation of iCA3 axons (vCA1d, 0.1 ± 0.04, n = 25 cells, N = 7 mice; vCA1m, 1.0 ± 0.2, n = 39 cells, N = 7 mice; vCA1v, 0.05 ± 0.03, n = 20 cells, N = 6 mice). Inset shows the example traces recorded from cells in vCA1d, vCA1m and vCA1v, respectively. Scale bars, 500 µm.

To confirm the dCA2/3→vCA1d pathway revealed by retrograde tracing, we performed anterograde tracing to visualize the brain-wide axon projection from dCA2/3, using local injection of GFP-expressing adeno-associated viral vectors (AAV) into dCA2/3. By measuring the axonal fluorescence intensities in vCA1 subregions, we found that the dCA2/3 axons indeed preferentially targeted vCA1d (Fig.1e and F; Fig. S3a). In contrast, the intermediate CA3 (iCA3) preferentially sent axons to vCA1m (Fig. 1g and h and Fig. S3b). These anterograde tracings together with retrograde tracings confirmed a preferential projection from dCA2/3 to vCA1d (Fig. 1i and j), which could be an anatomical route linking dorsal area with ventral area of hippocampus. Moreover, we visualized the axons from vCA1d neurons postsynaptic to dCA2/3 and observed prominent axonal arborizations in amygdala and prefrontal cortex (Fig. S4). These results suggest that the vCA1d is a key node to bridge the information flow from dorsal hippocampus to the amygdala.

The anatomical tracing results raised the possibility that dCA2/3 and iCA3 axons form distinct functional connections with vCA1d and vCA1m, respectively. To test this hypothesis, we performed patch-clamp recording in vCA1 subregions in acute brain slices with optogenetic stimulation of dCA2/3 axons after injecting AAV-CaMKIIα- channelrhodopsin-2 (ChR2) (Zhang et al., 2007) into dCA2/3. By recording the excitatory postsynaptic current (EPSC) in vCA1 cells, we found that dCA2/3 axons preferentially connected with vCA1d neurons (Fig. 1k-m), which was monosynaptic and glutamatergic (Fig S2e and f). In contrast, iCA3 axons preferentially connected with vCA1m cells (Fig. 1n-p). Taken together, these results demonstrate that dCA2/3 sends a specific functional projection to vCA1d along the longitudinal axis, in which vCA1d-projecting cells mainly locate in dCA2 and the distal part of dCA3.

### The dCA2/3**→**vCA1d pathway contributes to CFC

To determine whether the dCA2/3→vCA1d pathway is necessary for context-dependent associative learning, we performed optogenetic inhibition of dCA2/3→vCA1d axons in behaving mice during CFC. After bilateral injection of AAV-CaMKIIα-NpHR (Halorodopsin) (Gradinaru et al., 2010) into dCA2/3, the optical fibers were bilaterally implanted above vCA1d and NpHR-expressing axons were silenced by a yellow laser light (589 nm) when the foot shocks were delivered during CFC (Fig. 2a; Fig. S5a for fiber tip positions). During the fear retrieval in conditioned context, we found that conditioned freezing was significantly impaired by inhibiting dCA2/3→vCA1d axons during CFC (Fig. 2b and c; Fig. S6). On the contrary, inhibiting iCA3→vCA1m axons tended to increase the conditioned freezing, albeit not significantly (Fig. 2e and f; Fig. S5b). The generalized fear was neither affected by inhibition of dCA2/3→vCA1d pathway (Fig. 2d) nor by inhibition of iCA3→vCA1m pathway (Fig. 2g). We further examined the functional roles of these pathways in one-shock CFC (see Methods) by the same optogenetic manipulations during the foot shock (Fig. S7a). Optogenetic inhibitions of dCA2/3→vCA1d axons, but not iCA3→vCA1m axons, repeatedly impaired one-shock CFC (Fig. S7b and c).

**Figure 2.**
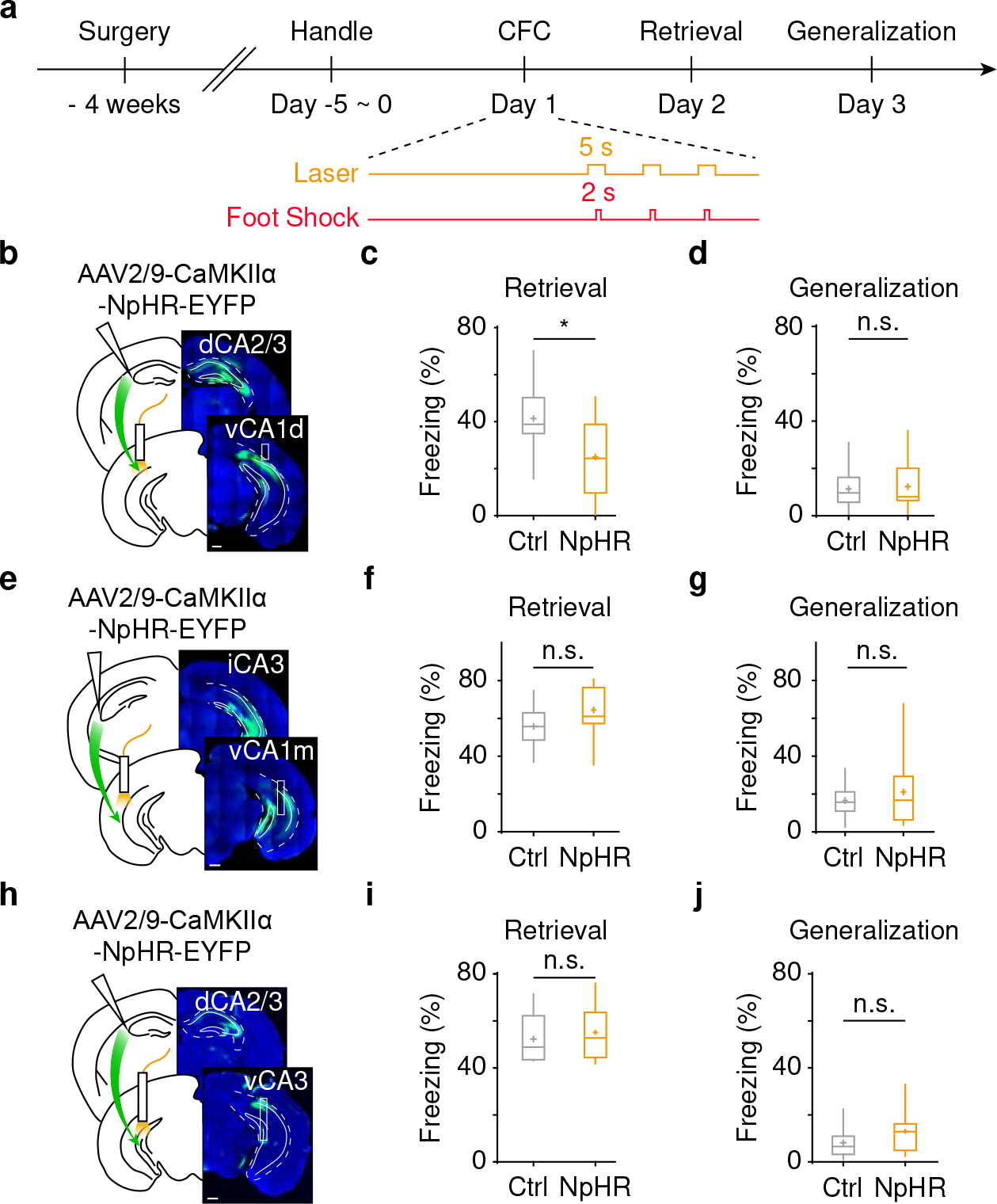
CFC requires dCA2/3**→**vCA1d pathway. (a) ) Scheme for experimental procedures and behavioral protocols with optogenetics. (b) Scheme showing AAV injections and fiber implantations and example pictures showing fluorescence of NpHR-EYFP in dCA2/3 and vCA1d, respectively. (c – d) Box plot of freezing in conditioned context (c, retrieval) and neutral context (d, generalization) after CFC. Conditioned freezing: 41.7 ± 3.5% Control (Ctrl) vs. 30.0 ± 3.4% NpHR; unpaired *t*-test, P = 0.024. Generalized freezing: 11.3 ± 2.1% Ctrl vs. 12.3 ± 2.7% NpHR; unpaired *t*-test, P = 0.77. N = 16 mice for Ctrl, N = 14 mice for NpHR. (e) Scheme showing AAV injections and fiber implantations and example pictures showing fluorescence of NpHR-EYFP in iCA3 and vCA1m, respectively. (f – g) Box plot of freezing in conditioned context (f, retrieval) and neutral context (g, generalization) after CFC. Conditioned freezing: 55.8 ± 2.9% Ctrl vs. 64.5 ± 3.6% NpHR; unpaired *t*-test, P = 0.072. Generalized freezing, 16.6 ± 2.4% Ctrl vs. 21.2 ± 5.1% NpHR; Mann Whitney U test, P = 0.89; N = 14 mice for Ctrl, N = 13 mice for NpHR. (h) Scheme showing AAV injections and fiber implantations and example pictures showing fluorescence of NpHR-EYFP in dCA2/3 and vCA3, respectively. (i – j) Box plot of freezing in conditioned context (i, retrieval) and neutral context (j, generalization) after CFC. Conditioned freezing: 52.3 ± 3.5% Ctrl vs. 55.1 ± 3.6% NpHR; unpaired *t*-test, P = 0.58. Generalized freezing: 8.2 ± 2.2% Ctrl vs. 13.0 ± 2.8% NpHR; Mann Whitney U test, P = 0.13. N = 9 mice for Ctrl, N = 10 mice for NpHR. Scale bars, 500 µm.

We next performed optogenetic manipulation of the activity of dCA2/3 axon terminals in ventral CA3 (vCA3), another downstream target of dCA2/3 in ventral hippocampus (Fig. 2h). We found that silencing dCA2/3→vCA3 axons did not affect conditioned fear (Fig. 2i-j, Fig.S5c). We further performed optogenetic manipulations of dCA2/3 axons to its downstream target in dorsal hippocampus, dorsal CA1 (dCA1).

Interestingly, we found that optogenetic inhibition of dCA2/3→dCA1 axons impaired the CFC (Fig. S7d-g).

Previous report has shown that dCA3 is necessary for novel context recognition (Wagatsuma et al., 2018). We thus inquired whether dCA2/3→vCA1d pathway is involved in contextual learning, in which animals show a high ratio of travelled distance in novel and recognized environmental contexts. When animals went through novel context recognition paradigm with optogenetic inhibition of dCA2/3→vCA1d axons, we observed significantly lower distance ratio in the NpHR group than in the control group (Fig. S8a- c). However, optogenetic inhibition of iCA3→vCA1m axons did not affect the distance ratio (Fig. S8d and e). Taken together, these results showed that the dCA2/3→vCA1d pathway represents a specific anatomical route from dorsal to ventral hippocampus, and demonstrate its specific and causal role in context recognition and contextual fear learning.

### Context-dependent activity of dCA2/3 and vCA1d neurons associated with CFC

Prior studies in rats demonstrated that the place cells in dCA3 (mostly in the proximal part) processed context information by context-dependent remapping of place fields (Leutgeb et al., 2007) and exhibited functional heterogeneities along the transverse axis (Lee et al., 2015; Lu et al., 2015). As the dCA2/3-to-vCA1d projection mostly originated from distal part of dCA3 and dCA2 (Fig. 1), we asked whether they are involved in context processing. We first confirmed that dCA2/3 was significantly activated by context exposure based on the cFos staining experiment (Fig. S9a-d). We then implanted tetrodes into dCA2/3 and performed single-unit recording when the animals were exploring two distinct contexts which were either morphed or separated (see Methods, Fig. S9e-q). We compared the neuronal activity of individual neurons when animals randomly explored two distinct contexts. We found that about 2/3 of recorded neurons exhibited context-modulated activities in both morphed and separated protocols (morphed, 58%; separated, 65%; Fig. S9e-m). The firing activity evoked by context exposures also differed in a context-specific manner for the whole neuronal population (Fig. S9i and n). Taken together, these results demonstrate that the neuronal activity in distal dCA2/3, an upstream of vCA1d, is prominently modulated by the environmental context.

We next sought to investigate the function role of dCA2/3 and vCA1d neurons in CFC by leveraging the head-mountable miniscope to perform deep-brain Ca^2+^ imaging in behaving mice(Cai et al., 2016). After injection of AAV-CaMKIIα-GCaMP6f into dCA2/3 or vCA1d and implantation with the gradient-index (GRIN) lenses, we performed pyramidal cell-specific Ca^2+^ imaging with single-cell resolution. We found that both dCA2/3 and vCA1d exhibited context-specific activities during the exposure to context A and B (Fig. S10), which was consistent with the results from single-unit recording (Fig. S9e-n). We then tracked the Ca^2+^ signals from the same population of cells when animals went through the CFC paradigm, in which animals were exposed to both context A and B before and after CFC in context A (Fig. 3a-e). During the CFC, we observed intermingled distributions of shock-responsive and shock non-responsive cells (SR and SNR cells) and significant larger proportion of SR cells in vCA1d than in dCA2/3 (Fig. 3f-h). This is consistent with photometric recording that foot shocks evoked more sensitized Ca^2+^ signals in vCA1d than in dCA2/3 (Fig. S11). Notably, the amygdala exhibited significantly larger Ca^2+^ responses to foot shocks than the dCA2/3 (Fig. S11a-d).

**Figure 3.**
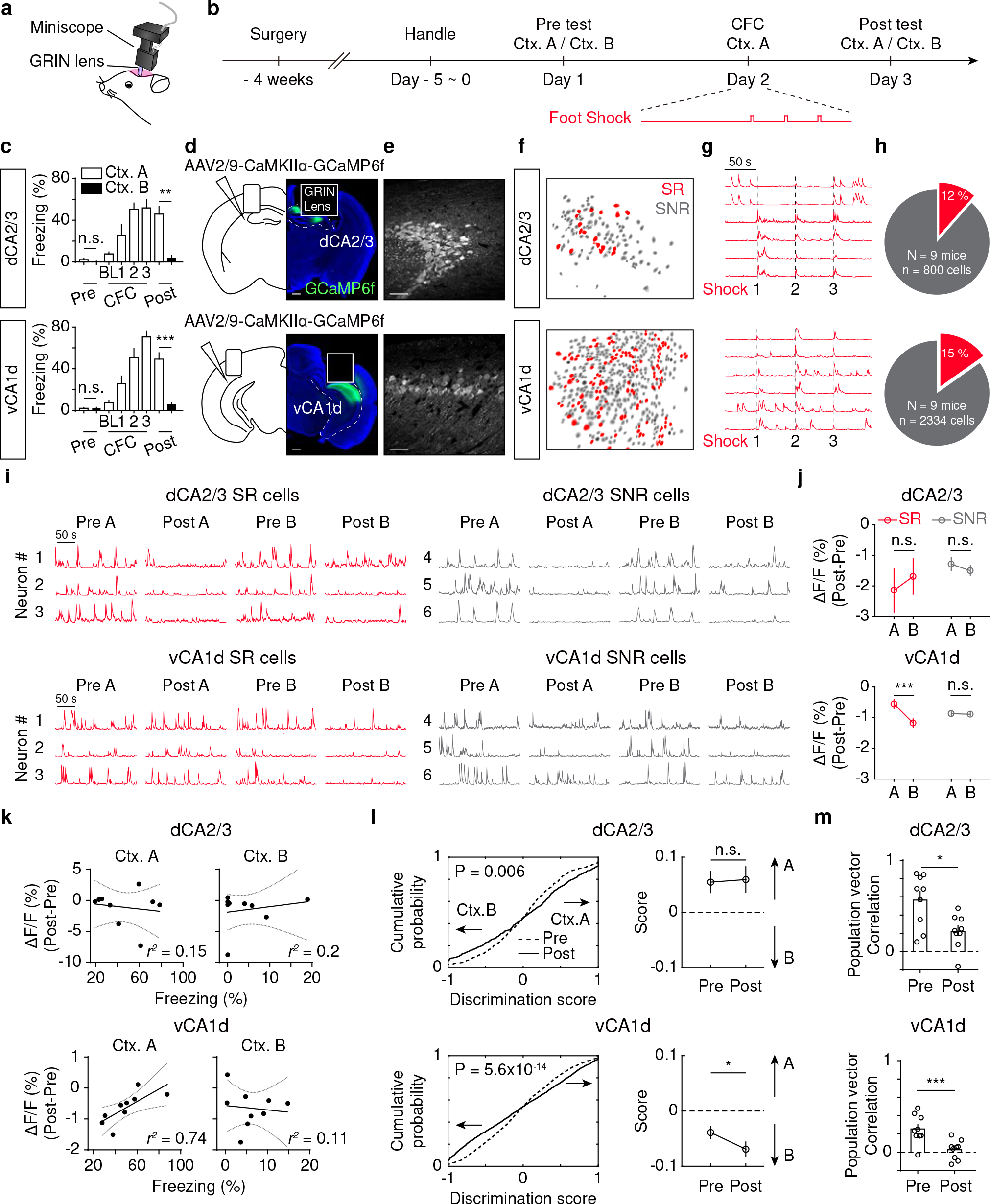
The post-CFC activity changes are context specific in vCA1d but not in dCA2/3. (a) ) Scheme for head-mountable miniscope with implanted GRIN lens. (b) Scheme for experimental procedures and behavioral protocols. (c) Summary of freezing in two distinct contexts before (Pre), during (CFC) and after (Post) the conditioning. Baseline (BL) refers the 3 min before 1^st^ shock. The freezing is significantly higher in the conditioned context (Ctx. A, open) than in the neutral context (Ctx. B, filled) after CFC. The dCA2/3 implantations, 45.9 ± 7.5% Ctx. A vs. 4.3 ± 2.1% Ctx. B; Wilcoxon signed rank test, P = 0.0039; N = 9 mice. The vCA1d implantations, 49.6 ± 6.0% Ctx. A vs. 6.1 ± 1.6% Ctx. B; paired *t*-test, P = 0.0002; N = 9 mice). (d) Scheme showing AAV injections and implantations of GRIN lens (left) and example pictures (right) showing fluorescence of GCaMP6f in dCA2/3 and vCA1d, respectively. (e) Example showing confocal images of GCaMP6f expression in dCA2/3 and vCA1d. (f) The cell map illustrating the spatial locations of shock-responsive (SR, red) and shock- non-responsive (SNR, grey) cells. (g) Example traces showing the Ca^2+^ signals upon foot shocks (lines). (h) Pie chart summary for the percentages of SR cells (shown in red: dCA2/3, 11.5%, n = 800 neurons from N = 9 mice; vCA1d, 15.2%, n = 2334 neurons from N = 9 mice, Chi- square test with Yates’ correction, P = 0.011). (i) Example traces showing the Ca^2+^ activity of SR (red) and SNR (grey) cells in context A and B before and after conditioning in A. (j) Summary of post-CFC changes in Ca^2+^ signals of SR and SNR cells as shown in (i). Ctx. A vs. Ctx. B, paired *t*-test: dCA2/3 SR cells, -2.14 ± 0.72% vs. -1.69 ± 0.59%, P = 0.40; dCA2/3 SNR cells, -1.29 ± 0.23% vs. -1.50 ± 0.18%, P = 0.22; vCA1d SR cells, - 0.55 ± 0.17% vs. -1.18 ± 0.15%, P = 9.70×10^-4^; vCA1d SNR cells, -0.87 ± 0.07% vs. -0.89 ± 0.06%, P = 0.81. (k) The correlation between changes of Ca^2+^ signal in Ctx. A and Ctx B versus contextual fear retrieval in individual animals (linear regression analysis with 95% confidence bands; dCA2/3: Ctx. A, *r^2^* = 0.15, P = 0.69; Ctx. B, *r^2^* = 0.2, P = 0.61; vCA1d: Ctx. A, *r^2^* = 0.74, P = 0.023; Ctx. B, *r^2^* = 0.11, P = 0.78). vCA1d cells before and after CFC (Kolmogorov-Smirnov test: dCA2/3, P = 0.006; vCA1d, P = 5.6×10^-14^). Right, the summary of context discrimination scores in Pre and Post (Pre vs. Post, paired *t*-test: dCA2/3, 0.05 ± 0.02 vs. 0.06 ± 0.02, P = 0.87; vCA1d, -0.04 ± 0.01 vs. -0.07 ± 0.01, P = 0.016). (m) Summary of population vector correlation of neuronal activity between context A and B in mice with CFC (Pre vs. Post, paired *t*-test; dCA2/3, 0.56 ± 0.1 vs. 0.22 ± 0.06, P = 0.022; vCA1d, 0.25 ± 0.05 vs. 0.03 ± 0.04, P = 0.0003). Data is summarized as mean ± SEM. Scale bars, 500 µm and 50 µm.

After fear conditioning, all animals with GRIN-lens implantation showed significantly higher freezing in the conditioned context (Ctx. A) than in the neutral context (Ctx. B) during the memory retrieval (Fig. 3c). Intriguingly, the Ca^2+^ signals were largely suppressed during the freezing episodes compared to non-freezing episodes in both dCA2/3 and vCA1d (Fig. S12). These results suggest that aversive stimuli evoke stronger Ca^2+^ signals in vCA1d than in dCA2/3, and that neuronal activity in both regions is modulated by the emotional state.

Next, we analyzed the changes of Ca^2+^ activities evoked by the same contexts before and after CFC. While overall Ca^2+^ signals were reduced after CFC, we noticed that vCA1d neurons showed a significantly smaller reduction than dCA2/3 neurons in both conditioned context A and neutral context B (Fig. S13a, see the control in Fig. S13c and d). Interestingly, we found that the SR cells in vCA1d maintained significantly higher Ca^2+^ activities during the contextual fear retrieval in conditioned context A than in neutral context B; whereas the SNR cells in vCA1d showed similar post-CFC changes of Ca^2+^ activities in both contexts (Fig. 3i and j). In contrast, neither SR nor SNR cells in dCA2/3 showed any context-specific changes of Ca^2+^ activities (Fig. 3i and j). These results showed that vCA1d neurons, but not dCA2/3 neurons, exhibited context-specific activity changes depending on US responses during CFC. At the behavioral level, the changes of vCA1d Ca^2+^ activities in conditioned context significantly correlated with the fear retrieval following CFC across individual animals (Fig. 3k). In contrast, no correlation was observed in dCA2/3 neurons (Fig. 3k). Taken together, these results indicate that the US signals instructs context- specific changes of Ca^2+^ activities in vCA1d neurons but not in dCA2/3 neurons.

We further assessed the discrimination of dCA2/3 and vCA1d neuronal activity in conditioned vs. neutral contexts by computing the context discrimination scores at single- cell level (see Methods). We found that the discrimination scores in both dCA2/3 and vCA1d neurons were significantly sharpened after CFC (Fig. 3l). Accordingly, the absolute discrimination scores were significantly enhanced (Fig. S13b). In the context re-exposures without CFC, the absolute discrimination scores were not changed at all in dCA2/3 neurons but were significantly increased in a small degree in vCA1d neurons (Fig. S13e). Notably, the net discrimination scores after CFC were biased towards the neutral context over the conditioned context in vCA1d neurons, but not in dCA2/3 neurons (Fig. 3l).

Finally, we assessed the populational neuronal activity in context discrimination by computing the population vector (PoV) correlation between conditioned and neutral contexts (see Methods). We found that the PoV correlation between two contexts were significantly decreased in both dCA2/3 and vCA1d neurons after CFC (Fig. 3m). This was not seen in the context re-exposures without CFC (Fig. S13f). Taken together, the Ca^2+^ imaging results with single-cell resolution demonstrate that only vCA1d pyramidal cells exhibited context-specific neuronal activity changes after CFC while both dCA2/3 and vCA1d neuronal activity showed sharpened discriminations between conditioned and neutral contexts.

### Enhanced monosynaptic excitation of pyramidal cells in vCA1d after CFC

The synaptic plasticity is fundamental for associative fear learning and memory (Fanselow and Poulos, 2005; LeDoux, 2000; Maren, 2001). The fact that vCA1d neurons showed a significantly smaller reduction of Ca^2+^ signals after CFC than dCA2/3 neurons promoted us to examine the synaptic plasticity of dCA2/3→vCA1d pathway upon CFC. We combined photometry and optogenetics in behaving animals to address whether dCA2/3→vCA1d pathway is potentiated *in vivo* after CFC. We injected AAV-CaMKIIα- GCaMP6f into vCA1d and AAV-CaMKIIα-ChR2 into dCA2/3 and implanted optic fiber above vCA1d (Fig. 4a-c). By measuring the vCA1d Ca^2+^ signals in response to optogenetic test stimulation of dCA2/3 neurons (see Methods), we monitored the dynamics of these Ca^2+^ signals as a proxy for projection strength of dCA2/3→vCA1d pathway. We found that Ca^2+^ signals in vCA1d pyramidal cells were significantly enhanced during the first 10 min after CFC, whereas no detectable changes were observed in control animals (Fig. 4d-g). To further address whether the similar potentiation happens in vCA1d neurons with direct dCA2/3 inputs, we injected AAV1-Cre and AAV-CaMKIIα-ChR2 into dCA2/3 and AAV- DIO-GCaMP6s into vCA1d (Fig. 4h and i). By doing so, we selectively recorded Ca^2+^ signals from vCA1d neurons that received direct projection from dCA2/3 in animals after CFC (Fig. 4h and i). Consistently, we observed significant increase of the Ca^2+^ signals in vCA1d neurons in response to optogenetic test stimulation of dCA2/3 neurons during fear retrieval on the next day after an overnight memory consolidation (Fig. 4j and k).

**Figure 4.**
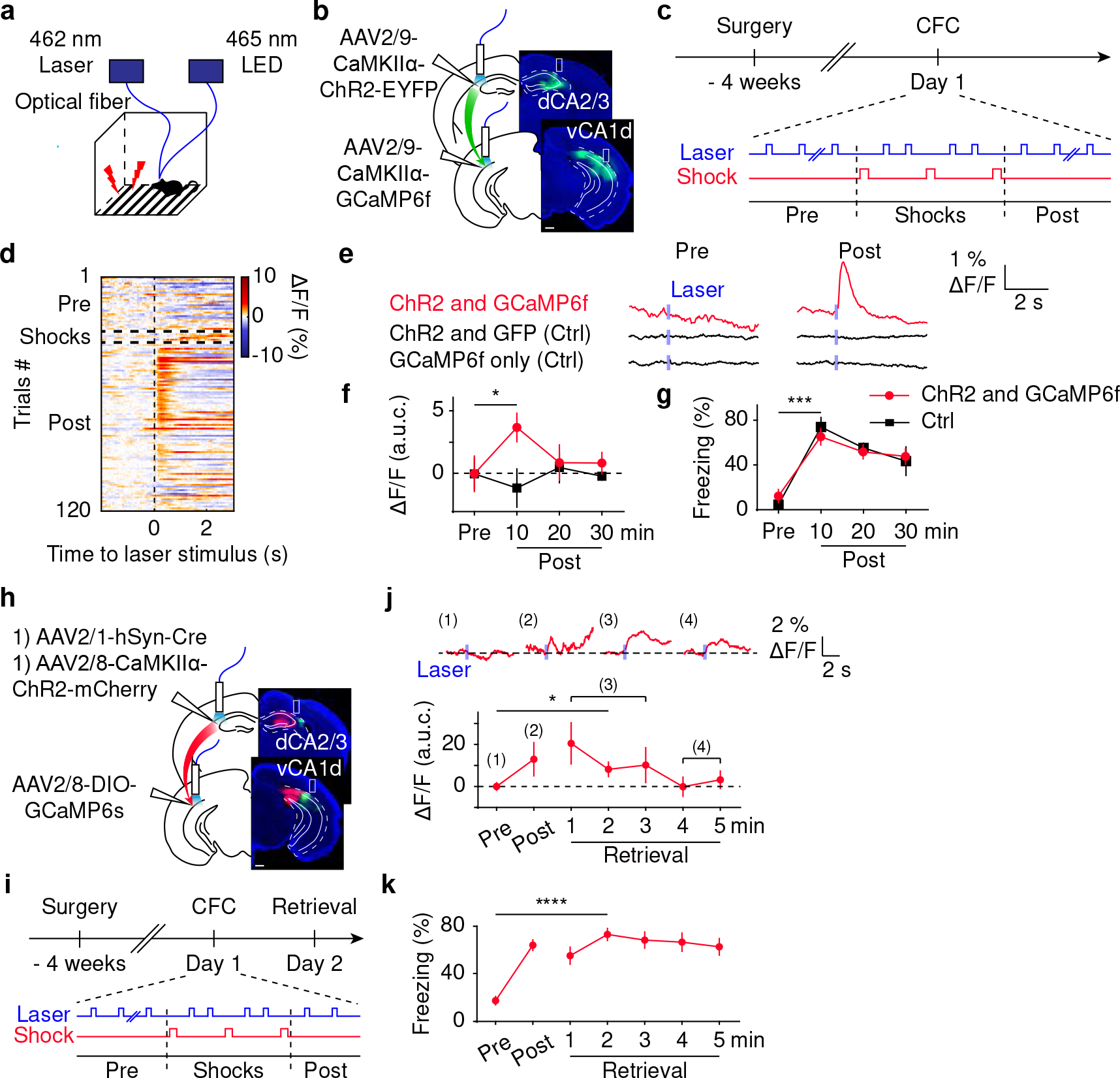
The dCA2/3**→**vCA1d projection onto pyramidal cells is potentiated *in vivo* by CFC. (a) Scheme illustrating combination of photometric recording and optogenetic testing pulses in animals going through CFC. (b) Scheme (left) illustrating AAV injections and fiber implantations and example pictures (right) showing fluorescence in dCA2/3 and vCA1d, respectively. (c) Scheme illustrating experimental procedures. The dCA2/3→vCA1d connection strength onto pyramidal cells was measured by the optogenetic testing pulses delivered to dCA2/3 at 0.05 Hz while photometry for vCA1d pyramidal cells was constantly on. Three shocks were delivered after 10 min baseline period (Pre) and followed by 30 min recording (Post). (d) Example heatmap showing Ca^2+^ signals for 120 trials recorded from vCA1d with optogenetic testing pulses. (e) Example photometric traces averaged from 10 min trials in Pre and Post periods, respectively, in animals going through the same behavioral protocol depicted in (c). The red example (top) had the same injection depicted in (b). The control groups were either injected with AAV-ChR2 in dCA2/3 and AAV-GFP in vCA1d (middle) or injected with only AAV-GCaMP6f in vCA1d (bottom). (f) ) Summary for Ca^2+^ signals during the Pre (10 min) and Post (30 min) periods (offset by subtracting the mean activity during the Pre period). The Ca^2+^ signals significantly increased in the first 10 min after CFC (3.7 ± 1.1, area under the curve [a.u.c.]; paired *t*- test, P = 0.031; N = 10 mice, red). No changes were observed in control animals (-1.1 ± 1.5 a.u.c.; paired *t*-test, P = 0.39; N = 4 mice, black). (g) Summary for freezing during fear conditioning depicted in (c). Freezing for the first 10 min after CFC: ChR2 & GCaMP6f group, 65.2 ± 7.3%, N = 10 mice, paired *t*-test, P = 0.0001; Ctrl group, 73.7 ± 8.5%; N = 4 mice, paired *t*-test, P = 0.0039. (h) Scheme (left) illustrating AAV injections and fiber implantations and example pictures (right) showing fluorescence in dCA2/3 and vCA1d, respectively. (i) Scheme illustrating experimental procedures to measure the dCA2/3→vCA1d connection strength onto pyramidal cells with direct inputs from dCA2/3. The behavioral protocol is the same as depicted in (c), except that the Pre period is 3 min and the Post period is only 1 min after CFC. The fear retrieval (5 min) is 1 day after CFC. (j) Top, example photometric traces averaged from trials in Pre (1), Post (2) and fear retrieval periods (3 and 4), respectively, in the animal going through the protocol depicted in (i). Bottom, summary for Ca^2+^ signals during the Pre (3 min) and Post (1 min) periods and fear retrieval (5 min) 1 day later (offset by subtracting the mean activity during the Pre period). The Ca^2+^ signals tend to increase during the fear retrieval 1 day after CFC (Retrieval at 2 min, 8.2 ± 3.8 a.u.c., Wilcoxon signed-rank test, P = 0.042, N = 11 mice). (k) Summary for freezing during CFC and retrieval depicted in (i). Pre vs. Retrieval at 2 min; 17.1 ± 4.1% vs. 73.1 ± 5.7%; N = 11 mice; paired *t*-test, P = 3.82×10^-5^. Data is summarized as mean ± SEM. Scale bars, 500 µm.

To further characterize the cellular mechanism of synaptic plasticity, we injected AAV-CaMKIIα-ChR2 into dCA2/3 and subjected animals to CFC. After the fear retrieval, we obtained acute brain slices of vCA1 and performed patch-clamp recording from vCA1d pyramidal cells in combination of optogenetic stimulation of ChR2-expressing dCA2/3 axons (Fig. 5a-c). We found that vCA1 pyramidal cells responded with larger EPSPs (excitatory postsynaptic potentials) and became more excitable after fear learning in CFC group than in control group without foot shocks (Fig. 5d and e). These results confirmed that the dCA2/3→vCA1d projection is strengthened after CFC.

**Figure 5.**
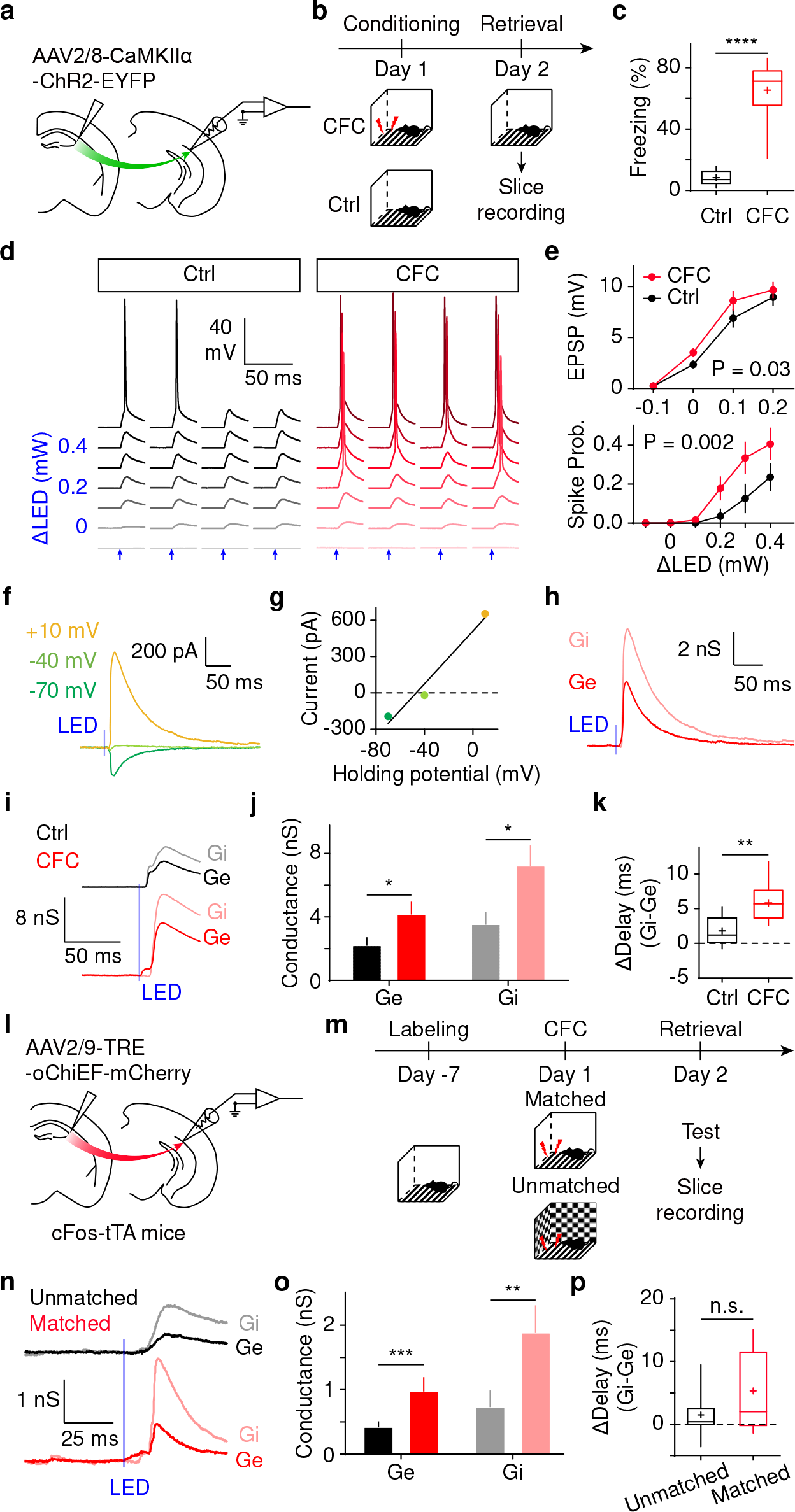
The excitability and synaptic conductances of vCA1d pyramidal cells are enhanced by CFC. (a and b) Schemes illustrating the AAV injection and behavioral protocols for CFC and Control (Ctrl) group. (c) Box plot for freezing during contextual fear retrieval. Ctrl, 8.4 ± 1.7%; N = 9 mice. CFC, 65.4 ± 6.8%; N = 9 mice. Unpaired *t*-test, P = 4.69×10^-7^. (d) Example showing whole-cell recordings of two vCA1d neurons from Ctrl (black) and CFC (red) groups, respectively, while blue LED stimuli (arrows) were delivered with increasing intensities (offset by minimal stimuli). (e) Summary of EPSP and spike probabilities evoked by the LED stimuli. The Ctrl vs. CFC: n = 26 cells, N = 7 mice for Ctrl, n = 29 cells, N = 7 mice for CFC (EPSP, two-way ANOVA, P = 0.03; Spike probability, two-way ANOVA, P = 0.002). (f) Example showing voltage-clamp whole-cell recordings of LED-evoked postsynaptic currents at holding potentials of -70, -40 and +10 mV. (g) The total synaptic conductance and reversal potential obtained by linear regression of synaptic currents and holding potentials. (h) Isolated excitatory and inhibitory conductances based on the total synaptic conductance and reversal potential obtained in (g). (i) Example traces of *Ge* and *Gi* upon LED stimuli (blue line) from two vCA1d neurons in Ctrl and CFC groups respectively. (j) Summary of *Ge* and *Gi* for vCA1d neurons from Ctrl and CFC groups. The Ctrl vs. CFC, unpaired *t*-test, Ctrl, n = 9 cells, N = 4 mice; CFC, n = 11 cells, N = 4 mice (*Ge*, 2.2 ± 0.4 nS Ctrl vs. 4.2 ± 0.7 nS CFC, P = 0.043; *Gi*: 3.5 ± 0.8 nS Ctrl vs. 7.2 ± 1.2 nS CFC, P = 0.025). (k) Box plot of delay differences between *Ge* and *Gi* for vCA1d neurons (same cells in [j]) from Ctrl and CFC groups (Unpaired *t*-test, Ctrl vs. CFC: 1.8 ± 0.7 ms vs. 5.8 ± 0.8 ms; P = 0.0021). (l) Scheme for AAV injection in cFos-tTA mice. (m) Scheme for experimental procedures with timelines. (n) Example traces of *Ge* and *Gi* in two vCA1d neurons from unmatched group (black, CFC in a different context instead of tagged context) and matched group (red, CFC in the tagged context). (o) Summary of *Ge* and *Gi* amplitudes for vCA1d neurons from matched and unmatched groups (Mann Whitney U test, unmatched vs. matched: *Ge*, 0.4 ± 0.1 nS vs. 1.0 ± 0.2 nS, P = 0.0009; *Gi*, 0.7 ± 0.2 nS vs. 1.9 ± 0.4 nS, P = 0.0052; n = 20 cells, N = 5 mice for unmatched group; n = 16 cells, N = 5 mice for matched group). (p) Box plot of delay differences between *Ge* and *Gi* for vCA1d neurons from matched and unmatched groups (unmatched, 1.5 ± 1.0 ms, n = 12 cells; matched, 5.3 ± 1.8 ms, n = 13 cells; unpaired *t*-test, P = 0.089). Data is summarized as mean ± SEM.

To dissect the synaptic mechanism underlying the potentiation of dCA2/3→vCA1d pathway, we isolated the excitatory and inhibitory synaptic conductances (*Ge* and *Gi*) from synaptic currents in vCA1d neurons evoked by optogenetic stimulation of dCA2/3 axons at varying holding potentials (see Methods, Fig. 5f-h). We found that the amplitudes of both *Ge* and *Gi* were enhanced in the CFC group compared to the control group (Fig. 5i and j). Interestingly, the delay differences between *Ge* and *Gi* became significantly larger in the CFC group (Fig. 5k). These results suggest that the CFC enhances amplitudes of both *Ge* and *Gi* onto vCA1d neurons, in which *Gi* rises much more slowly than *Ge*.

### Context-specific potentiation of the dCA2/3**→**vCA1d projection after CFC

To further investigate whether the strengthening of dCA2/3→vCA1d pathway is context specific, we injected AAV-TRE-oChiEF (Lin et al., 2009) into dCA2/3 in *cFos-tTA* mice (Reijmers et al., 2007). Animals were randomly assigned into two groups and all were exposed to context A. The dCA2/3 neurons activated during the context exposure (in the absence of doxycycline) were tagged by oChiEF under the control of the cFos promoter and a tetracycline-controlled transactivator (tTA). One group went through CFC in context A (same as tagged context, matched group) and the other in context B (unmatched group). After fear retrieval following CFC, we obtained hippocampal brain slices and performed patch-clamp recording from vCA1d pyramidal cells while optogenetically stimulating oChiEF-expressing axons from tagged dCA2/3 neurons (Fig. 5l and m). We found that the amplitudes of *Ge* and *Gi* were significantly larger in the matched group than in the unmatched group (Fig. 5n and o). Furthermore, the delay differences between *Ge* and *Gi* tended to be larger, albeit not significantly, in the matched group than in the unmatched group (unpaired *t*-test, P = 0.089; Fig. 5p). Taken together, these results suggest that CFC potentiated dCA2/3→vCA1d pathway in a context-specific manner, via concerted changes in amplitudes and delays of *Ge* and *Gi*onto vCA1d pyramidal cells.

### Reduced SOM-cell-mediated disynaptic inhibition in vCA1d after CFC

The prominent *Gi* recorded from vCA1d pyramidal cells indicated that they received strong feed-forward inhibition mediated by local interneurons. To investigate whether and how subtype of interneurons in vCA1d receives inputs from dCA2/3, we leveraged the rabies vectors (Wickersham et al., 2007) to perform cell-type specific retrograde transsynaptic tracing from PV (parvalbumin), SOM, VIP (vasoactive intestinal peptide) interneurons and PN (pyramidal neurons) in vCA1d (Fig. 6a and b). We found that all interneuron subtypes received prominent inputs from dCA2/3 (Fig. 6c and d). Within hippocampus, interneurons received significantly higher proportions of inputs than PN (Fig. S14). Interestingly, while PN received significant larger proportion of inputs from sensory cortex than all three major subtypes of interneurons, SOM cells received significantly more inputs from sensory cortex than VIP cells (Fig. S14b). These results support the notion that all these types of vCA1d interneurons are likely to mediate feed-forward inhibition to vCA1d pyramidal cells driven by dCA2/3 projection.

**Figure 6.**
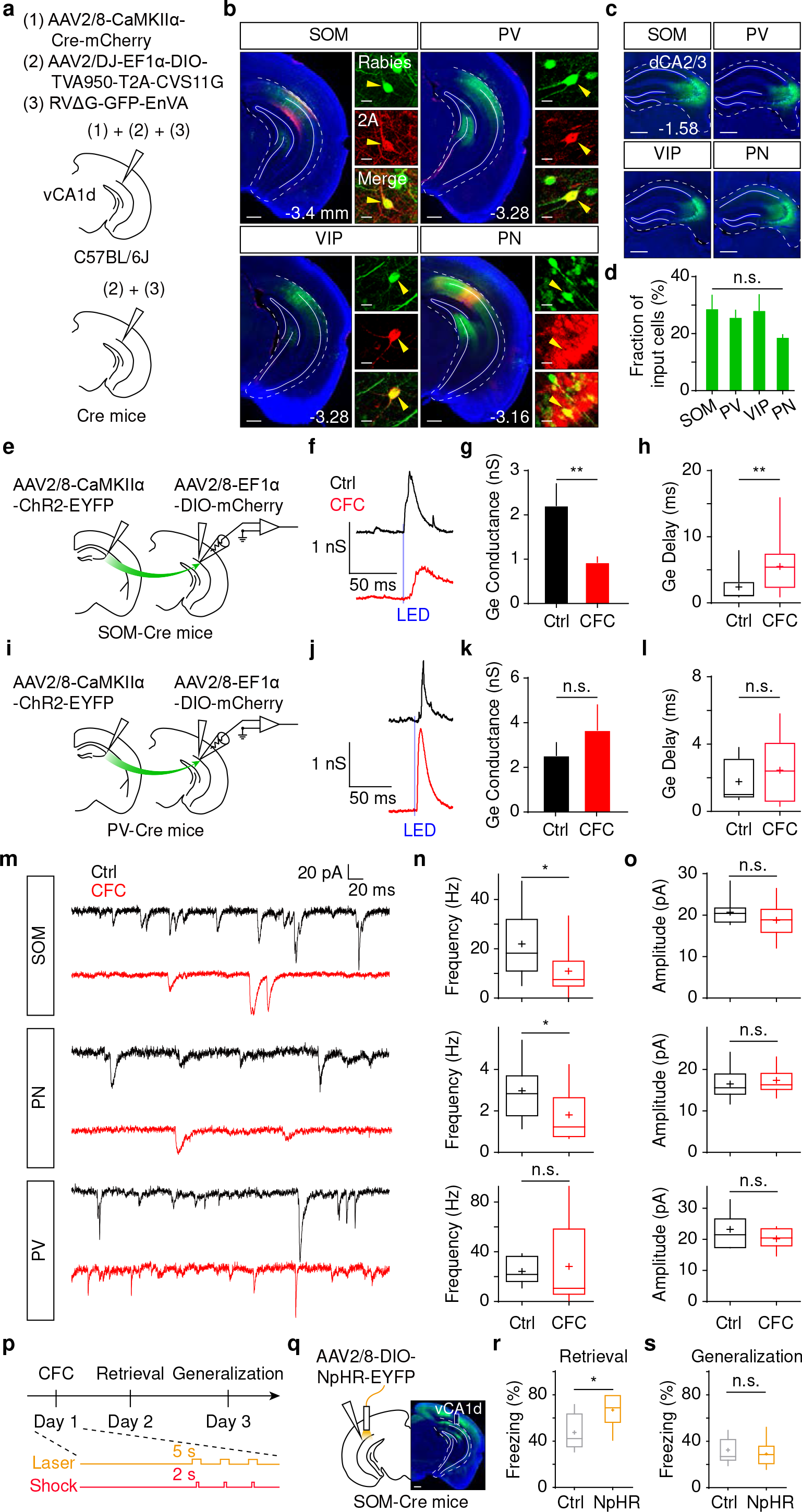
CFC-induced changes of synaptic inputs onto interneuron subtypes in vCA1d. (a) ) Scheme for AAV injections in C57BL/6J mice and Cre-expressing mice. (b) Examples showing fluorescence in the region of starter cells. Rabies-labeled cells are shown in green. 2A signals are shown in red. (c) Examples showing rabies-labeled cells in dCA2/3. (d) Summary of proportion of presynaptic cells in dCA2/3. One-way ANOVA, F(3,8) = 1.168, P = 0.38. (e and i) Scheme for AAV injections in SOM-Cre and PV-Cre mice. (f and j) Example traces of *Ge* and *Gi* in SOM and PV cells in vCA1d from Ctrl (black) and CFC (red) groups. (g and k) Summary (mean ± SEM) of *Ge* amplitudes in SOM and PV cells in vCA1d. SOM cells: Mann-Whitney U test, Ctrl vs. CFC: 2.2 ± 0.5 nS vs. 0.9 ± 0.1 nS; P = 0.0091; n = 13 cells, N = 3 mice for Ctrl; n = 17 cells, N = 5 mice for CFC. PV cells: Mann Whitney U test, Ctrl vs. CFC: 2.5 ± 0.6 nS vs. 3.7 ± 1.1 nS; P = 0.48; n = 13 cells, N = 4 mice for Ctrl; n = 10 cells, N = 4 mice for CFC). (h and l) Box plot of the *Ge* delay in SOM and PV cells in vCA1d. SOM cells: Mann- Whitney U test, Ctrl vs. CFC: 2.3 ± 0.6 ms vs. 5.4 ± 0.9 ms; P = 0.0062; PV cells: Unpaired *t*-test, Ctrl vs. CFC: 1.8 ± 0.3 nS vs. 2.5 ± 0.6 nS; P = 0.32. (m) Example traces of sEPSC in SOM, PN and PV cells in vCA1d in Ctrl (black) and CFC (red) groups. Voltage-clamp holding potential is -70 mV. (n and o) Box plot of sEPSC frequencies (n) and amplitudes (o) in SOM, PN and PV cells from Ctrl and CFC animals, respectively. SOM cells: 22.0 ± 4.6 Hz Ctrl vs. 10.9 ± 2.4 Hz CFC; Mann-Whitney U test, P = 0.02; n = 9 cells, N = 3 mice for Ctrl; n = 16 cells, N = 5 mice for CFC. PN cells: 3.0 ± 0.4 Hz Ctrl vs. 1.8 ± 0.4 Hz CFC; unpaired *t*-test, P = 0.044; n = 12 cells, N = 4 mice for Ctrl; n = 11 cells, N = 4 mice for CFC. PV cells: vs. 24.2 ± 2.9 Hz Ctrl vs. 28.2 ± 10.4 Hz CFC; unpaired *t*-test, P = 0.70; n = 11 cells, N = 5 mice for Ctrl; n = 10 cells, N = 5 mice for CFC. (p) Scheme illustrating experimental procedures. (q) Scheme showing AAV injections and fiber implantations and example pictures showing fluorescence of NpHR-EYFP in vCA1d of SOM-Cre mice. (r and s) Box plot of freezing in fear retrieval (r) and fear generalization (s) after CFC (conditioned freezing, 47.5 ± 5.0% Ctrl vs. 66.9 ± 5.4% NpHR; unpaired *t*-test, P = 0.02; generalized freezing, 32.5 ± 3.6% Ctrl vs. 29.1 ± 4.6% NpHR; unpaired *t*-test, P = 0.57; N = 9 mice for Ctrl, N = 7 mice for NpHR). Scale bars, 500 and 20 µm.

Next, we investigated the synaptic plasticity of dCA2/3 inputs onto interneuron subtypes which may govern the feed-forward inhibition onto pyramidal cells in vCA1d. We performed patch-clamp recording from SOM and PV cells in Cre mice where interneuron subtypes were labeled by AAV-DIO-mCherry and dCA2/3 axons were infected by ChR2-expressing AAV (Fig. 6e and i). While *Gi* was hardly detectable in SOM cells, we found that the *Ge* amplitude was significantly reduced (Fig. 6f and g) and the *Ge* delay was significantly increased in SOM cells after fear learning (Fig. 6f and h). However, these kinds of changes in *Ge* were not seen in PV cells (Fig. 6j-l). Notably, we observed a significant reduction of the frequency of spontaneous EPSCs in SOM cells after CFC (Fig. 6m-o), indicating a global reduction of presynaptic inputs. Similar reduction was observed in pyramidal cells but not in PV cells (Fig. 6m-o). These results suggest that CFC weakened synaptic inputs from dCA2/3 to SOM cells and thereby delayed feed-forward inhibition onto pyramidal cells in vCA1d.

The potential functional role of SOM-cell-mediated feed-forward inhibition during CFC was highlighted by the fact that vCA1d SOM cells received reduced *Ge* from dCA2/3 and decreased frequency of spontaneous EPSCs after CFC (Fig. 6f-h and m-o). Accordingly, inhibiting SOM cells may relive the feed-forward inhibition onto pyramidal cells in vCA1 and further boost the CFC. To test this hypothesis, we injected AAV-EF1α- DIO-NpHR into vCA1d of *SOM-Cre* mice and performed optogenetic inhibition of SOM cells in vCA1d during CFC (Fig. 6p and q). We found that contextual fear retrieval was significantly enhanced in animals expressing NpHR compared to control animals injected with AAV-EF1α-DIO-mCherry (Fig. 6r). The generalized fear was not affected (Fig. 6s). Taken together, these results demonstrate that CFC is gated by vCA1d SOM cells, via controlling feed-forward inhibition driven by dCA2/3→vCA1d projection.

## Discussion

Contextual learning and memory is essential for the animal to choose appropriate behaviors depending on the environmental context. In this study, we have identified a specific intra- hippocampal projection from dCA2/3 to vCA1d that conveys context information. Miniscope Ca^2+^ imaging with single-cell resolution revealed that neuronal activities in dCA2/3 and vCA1d could code for context information, but conditioned fear retrieval only correlated with activity changes in vCA1d but not in dCA2/3. These results support the notion that dCA2/3 is only involved in the context processing per se, whereas vCA1d severs as a critical site for initiating context-dependent association with emotional experience. Our study also unveils an orchestrated synaptic plasticity of monosynaptic excitatory and feed-forward inhibitory circuit in vCA1d that underlies the context- dependent associative learning.

Our work uncovers an essential role of the vCA1 in integrating context information from dCA2/3 with emotional experience and in broadcasting the information to its downstream areas such as amygdala and prefrontal cortex, as shown by axon tracing of dCA2/3-targeted vCA1d neurons (Fig. S4). Anatomically, there are several possible routes from the dorsal hippocampus to vCA1, the major output of ventral hippocampus. We have focused on the direct projection from dCA2/3 to vCA1d in this study. Alternative routes include indirect projections of dCA2/3→vCA3→vCA1d and dCA2/3→dCA1→vCA1d. These alternative pathways are unlikely to be involved in CFC, since we found that CFC was not affected by optogenetic silencing of dCA2/3→vCA3 projections (Fig. 2h-j) and connectivity between dCA1 and vCA1 is relatively sparse (Yang et al., 2014).

Our physiological recording studies demonstrated a CFC-induced strengthening of dCA2/3→vCA1d projection, resulting from an increased amplitude of monosynaptic excitatory conductance *Ge* and a delayed onset of inhibitory conductance *Gi* in vCA1d pyramidal cells (Fig. 5i-k). The delayed onset of *Gi* in vCA1d neurons could be explained by the decreased and delayed *Ge* in SOM-expressing interneurons involved in disynaptic inhibition (Fig. 6e-h). On the other hand, the increased *Gi* amplitude in vCA1d neurons is likely resulted from CFC-induced synaptic modification in other types of interneurons (e.g. subgroup of PV cells, Fig. 6i-l). The enhanced but delayed feed-forward inhibition onto vCA1d pyramidal cells may generate a sparse-coding activity pattern for better discrimination of conditioned vs. neutral context. Such mechanism has been proposed for sparse-coding in thalamocortical circuits (Gabernet et al., 2005).

We observed a striking post-CFC reduction of vCA1d neuron activity (Fig. S13a), consistent with a previous report (Jimenez et al., 2020). This is not contradictory to the finding of CFC-induced strengthening of dCA2/3→vCA1d projection, because there was an even larger reduction of presynaptic dCA2/3 neuronal activity (Fig. S13a). Furthermore, reduced vCA1d neuron activity could result in part from the decrease in the frequency of spontaneous EPSCs in vCA1d cells associated with CFC-induced elevation of evoked EPSCs (Fig. 6m-o). The mechanism underlying CFC-induced reduction of dCA2/3 neuron activity remains to be future examined.

The unconditional stimulus (US) driving associative learning is known to evoke hippocampal activity through cholinergic projection from the medial septum(Lovett- Barron et al., 2014). We have observed strong US signals in vCA1d neurons, as shown by foot shock-induced neuronal responses in both dCA2/3 and vCA1d, with larger number of neurons responding in the latter (Fig. 3g and h). We show that after CFC neuronal activity in vCA1d SR (shock-responsive) neurons exhibited significantly higher activity in conditioned context than that in neutral context, and this was not observed in SNR (shock- nonresponsive) neurons (Fig. 3j). This indicates that the US signals were integrated with the context information primarily in SR neurons in vCA1d. Surprisingly, we found no context-dependent activity change in both SR and SNR neurons in dCA2/3 (Fig. 3j). This suggests that association of emotional experience occurred in vCA1d but not in dCA2/3. In support of this notion, we found that magnitude of the changes in conditioned context- evoked neuron activity in vCA1d strongly correlated with the fear retrieval at the level of individual animals, and such correlation was absent in dCA2/3 (Fig. 3k). Thus, we conclude that the dCA2/3 is more likely to be involved in processing context information, whereas vCA1d represent an early hippocampal site for initiating contextual association with emotional experience.

Previous reports have shown that the dCA3 is capable of processing context information (Leutgeb et al., 2007; Lu et al., 2015) and is necessary for context recognition (Wagatsuma et al., 2018) and contextual memory retrieval by means of pattern completion (Nakazawa et al., 2002). The dCA2 is known to be important for social memory (Hitti and Siegelbaum, 2014). Whether dCA2 and dCA3 projections to vCA1d have distinct functions remains to be clarified. Our work supports a working model that the environmental context information is actively processed in the dCA2/3 area and routed to the downstream vCA1d area for integration with the US signal, where associative information is broadcasted to a wide range of brain areas such as amygdala and prefrontal cortex. Our findings pave the way for further dissection of intra-hippocampal longitudinal circuits underlying associative learning and memory.

## Acknowledgments

We thank all members of the Xu lab, L. Xu, A. Lüthi and M.-M. Poo for helpful discussions and comments. We thank C.-Y. Li, L.-N. Lin, Y. Zhou, L.Ma and N.-L. Xu for technical support and sharing resources and thank B. Roska and E.M. Callaway for sharing rabies- related cell lines and plasmids. This study was supported by National Key R&D Program of China, the Strategic Priority Research Program of the Chinese Academy of Sciences (XDB32010105), Shanghai Municipal Science and Technology Major Project (2018SHZDZX05), the National Natural Science Foundation of China (31771180, 91732106) and Shanghai Science and Technology Committee (19490711800).

## Author contributions

H.-S.C., S.Q. and C.X. conceived the project. C.X. supervised the work in the study. R.-R.Y. genotyped the transgenic mice and generated the rabies vectors. H.-S.C., S.Q. and R.- R.Y. performed rabies-based tracing and analyzed the results. H.-S.C., S.Q., G.-L.W., R.-R.Y. and N.Z. performed the AAV injection, histology and immunostaining and analyzed the results. G.-L.W. performed the patch-clamp recording and analyzed the results. S.Q. and Q.-X.Y. performed the single-unit recording and analyzed the results. J.-N.W. performed the cFos experiments and analyzed the results. H.-S.C. performed the photometric recording and analyzed the results. H.-S.C., S.Q. N.Z. and C.X. performed the optogenetic experiments and miniscope imaging and analyzed the data. H.-S.C. and C.X. wrote the manuscript with inputs from all authors.

Declaration of interests

The authors declare no competing interests.

## Data availability

All data generated or analyzed during this study are included in this published article (and its supplementary information files). Source data are provided with this paper.

## Code availability

The codes used for analysis in this manuscript are available from the corresponding author (C.X.) upon reasonable request.

## Methods details

### Animals

Animals were housed under a 12 h light / dark cycle and provided with food and water *ad libitum* in the animal facility of Institute of Neuroscience (CAS Center for Excellence in Brain Science and Intelligence Technology). All animal procedures were performed in accordance with institutional guidelines and were approved by the Institutional Animal Care and Use Committee (IACUC) of the Institute of Neuroscience. Wild-type C57BL/6J (Slac Laboratory Animal, Shanghai), *cFos-tTA* (Reijmers et al., 2007), *PV-Cre* (Hippenmeyer et al., 2005), *VIP-Cre* and *SOM-Cre* (Taniguchi et al., 2011) mice were used. All of the experimental mice used in the study were adult (over 8 weeks) male mice.

### Viral vectors

The SADΔG rabies vectors were generated as described before(Xu et al., 2016). In brief, rabiesΔG-GFP vectors were amplified from viral stocks in B7GG cells (gift from Ed Callaway, Salk Institute). EnvA pseudotyped rabies was produced in BHK-EnVA cells, concentrated by two rounds of centrifugation, suspended in Hank’s Balanced Salt Solution (Gibco) and titered in HEK293-TVA cells (gift from J. A. T. Young, Salk Institute) with serial dilutions of the virus. The titers of the rabiesΔG-GFP-EnVA were in the range of 10^8^– 10^9^ infectious units/ml. Virus was stored at -80°C until further use.

The following AAV vectors were produced at gene editing facility at the Institute of Neuroscience (titers in genome copies/mL). AAV2/9-CaMKIIα-hChR2(H134R)-EYFP (4.0 × 10^13^), AAV2/8-CaMKIIα-hChR2(H134R)-EYFP (2.4 × 10^13^), AAV2/8-CaMKIIα-Cre-mCherry (3.6 × 10^13^), AAV2/8-hSyn-DIO-GCaMP6s (2.88 × 10^13^), AAV2/8-CaMKIIα-DIO-eNpHR3.0-EYFP (3.8 × 10^13^), AAV2/8-EF1α-DIO-mCherry (1.9 × 10^13^), AAV2/DJ-EF1α-DIO-TVA950-T2A-CVS11G (plasmid from B. Roska, FMI; 1.6 × 10^13^).

The following AAV vectors were purchased from WeiZhen Biosciences, JiNan. AAV2/8- hSyn-GFP (1.0 × 10^13^), AAV2/8-CaMKIIα-ChR2-mCherry (1.82 × 10^13^), AAV2/1-hSyn-Cre (1.0 × 10^13^), AAV2/9-CaMKIIα-EYFP (1.28 × 10^13^). The following were purchased from Taitool Bioscience, Shanghai, AAV2/9-TRE-oChiEF-mCherry (1.6 × 10^13^), AAV2/9- CaMKIIα-GCaMP6f (1.38 × 10^13^), AAV2/9-CaMKIIα-eNpHR3.0-EYFP (1.52 × 10^13^).

### Stereotaxic injections

Mice were anaesthetized by isoflurane (induction 5%, maintenance 2%, RWD R510IP, China) and fixed in a stereotactic frame (RWD, China). Local analgesics (Lidocaine, Shandong Hualu Pharmaceutical) were administered before the surgery. The body temperature was maintained at 35°C by a feedback-controlled heating pad (FHC Inc). The dye and virus solutions were loaded into glass pipettes (tip diameter 10 – 20 µm) connected to a Picospritzer III (Parker Hannifin Corporation) and injected at following coordinates (posterior to Bregma, AP; lateral to the midline, LAT; below the brain surface, DV; in mm): dorsal CA2/3: AP -1.5, LAT ± 2.1, DV -1.7; intermediate CA3: AP -2.3, LAT ±3.1, DV -2.6; ventral CA1: AP -3.2, LAT ± 3.4, DV -1.5 (dorsal part), 2.6 (median part) and 3.5 (ventral part); basolateral amygdala: AP -1.2, LAT ± 3.0, DV -4.35. The pipettes stayed at injection sites for at least 3 min after injection.

*Retrograde tracing by retrobeads:* 300 nL red Retrobeads (fluorophore-coated latex beads, Lumafluor R-180) were diluted with PBS at 1: 100 and injected into vCA1 subregions. One week later, brain slices were prepared from injected animals.

*Cell-type specific rabies tracing:* For retrograde tracing from interneurons, 200 nL of AAV- EF1a-DIO-TVA-2a-G were injected into vCA1d of transgenic Cre mice. For retrograde tracing from pyramidal cells, a mixture (200 nL, ratio 1:1) of AAV-EF1a-DIO-TVA-2a-G and AAV-CaMKIIα-Cre-mCherry were injected into vCA1d of C57 mice. Two weeks later, 300 nL of rabiesΔG-GFP-EnVA were injected into vCA1d. The histology was done one week later.

*Anterograde tracing:* 300 nL AAV-hSyn-GFP were injected into dCA2/3 or iCA3. The histology was done 4 weeks later.

*Slice electrophysiology:* For C57 mice, 300 nL AAV-CaMKIIα-ChR2-EYFP was injected into dCA2/3 or iCA3. The slice recording was performed 2 weeks after injection. For Cre mice, 300 nL AAV-CaMKIIα-ChR2-EYFP was injected into dCA2/3 and 300 nL AAV- DIO-mCherry was injected into vCA1d. The slice recording was performed 2 weeks after injection. Patch-clamp recording was performed on mCherry-positive cells. For *cFos-tTA* mice, 500 nL AAV-TRE-oChieF-mCherry was injected into dCA2/3.

*Photometry recording combined with optogenetics:* 300 nL AAV-CaMKIIα-ChR2-EYFP was injected into dCA2/3 and 300 nL AAV-CaMKIIα-GCaMP6f was injected into vCA1d. For control animals, AAV-CaMKIIα-ChR2-EYFP was omitted or AAV-CaMKIIα- GCaMP6f was replaced by AAV-hSyn-GFP. In another set of experiments in Fig. 4h, a mixture (300 nL, ratio 1:1) of AAV1-hSyn-Cre and AAV-CaMKIIα-ChR2-mCherry was injected into dCA2/3 and 300 nL AAV-DIO-GCaMP6s was injected into vCA1d.

*Multi-channel photometry recording:* The dCA2/3, vCA1d and basolateral amygdala in the same mice were injected with 300 nL AAV-CaMKIIα-GCaMP6f, respectively.

*Miniscope recording:* 400 nL AAV-CaMKIIα-GCaMP6f was injected into dCA2/3 or vCA1d. GRIN lens was implanted 4 weeks after injection. Miniscope recording was performed 6 – 8 weeks after injection.

*Optogenetics in behaving mice:* For C57 mice, 300 nL AAV-CaMKII-NpHR-EYFP was bilaterally injected into dCA2/3 or iCA3. Control animals were injected with AAV- CaMKIIα-EYFP instead. For SOM-Cre mice, 400 nL AAV-DIO-NpHR-EYFP was bilaterally injected into vCA1d. Control animals were injected with AAV-EF1α-DIO-mCherry instead.

### Histology and immunohistochemistry

One to four weeks after viral injections, mice were transcardially perfused with phosphate buffered saline (PBS) followed by 4% (w/v) paraformaldehyde (PFA) in PBS. Brains were post-fixed in PFA overnight at 4°C. and then cut into 80 µm thick coronal slices with a vibratome (Leica, Germany). For immunostaining of free-floating sections, sections were incubated in blocking solution (3% bovine serum albumin and 0.5% Triton X-100 in PBS) for 1 h at room temperature and then incubated in blocking solution containing mouse anti-NeuN (1:500, Merck, MAB377), rabbit anti-PCP4 (1:200, Sigma, HPA005792-100UL), rabbit anti-2A (1:500, Merck, ABS31) antibodies over night at 4°C. Subsequently, sections were washed with PBS three times (10 min each) and incubated in blocking solution for 2 h at room temperature with fluorescent donkey anti-rabbit alexa flour 594 (1:500, Invitrogen, A21207), goat anti-rabbit alexa flour 647 (1:500, Invitrogen, A21245), donkey anti-rabbit alexa flour 488 (1:500, Invitrogen, A21206), goat anti-mouse alexa flour 350 (1:500, Invitrogen, A21049). Finally, immuno-labelled sections were rinsed three times with PBS, mounted with an anti-fade mounting media (Solarbio, S2100) dehydrated and coverslipped.

*Quantification of axonal fluorescence:* Images from brain slices were acquired by an Olympus FV3000 confocal microscope with same settings. For analysis of endogenous fluorescence intensities, hippocampal contours were drawn in Fiji (ImageJ). Mean fluorescence value of hippocampal slices were measured after background subtraction. Four slices per brain were analyzed.

*3D reconstruction of anatomical tracing:* Images from brain slices with rabies tracing and retrobeads tracing were acquired by an Olympus VS120 fluorescence microscope or a Zeiss Axio Scan.Z1 and reconstructed as described before(Xu et al., 2016). In brief, TIFF images were sorted, aligned (Fiji TrackEM plugin) and imported into Imaris (v7.3, Bitplane) for cell counting. For retrobeads tracing, the 3D distributions of counted cells exported by the Imaris XTensions (7.3.1) and analyzed in Matlab (MathWorks). These normalized cell numbers were represented by color-coded density map (200 µm square window, 100 µm sliding step). For rabies tracing, the brain slices with starter cells were imaged with confocal microscope and analyzed for co-localization between 2a staining and rabies fluorescence. For each brain sample, the cells that were not co-labeled with 2a fluorescence were identified as presynaptic input cells. The cell number for individual brain regions or slice sections was normalized by the total number of input cells in the whole brain.

*cFos mapping*: C57BL6/J mice were singly housed and habituated to the handling and transportation to the behavioral room for four days. On the experiment day, mice rested in the home cage for one hour after arrival in the behavioral room to minimize the background of cFos activity. Then mice were taken out from their homecage and placed into either a neutral context (in cm, 30 [L] × 30 [W] × 35 [H]) or the homecage for 10 min. Next, mice rested in the homecage for 90 min and were perfused with 4% PFA. Following 24 h post-fixation, free-floating 80-μm coronal sections were prepared using a vibratome (Leica) and incubated in the blocking solution (5% BSA, 0.5% Triton X in PBS) for 1 h at room temperature. The sections were then incubated with the anti-cFos rabbit primary antibody (SYSY; one batch with 1:10000 and the other batch with 1:5000; both batches yielded similar results and were pool together) for 18 h at 4 °C. Next, sections were washed with PBS and incubated with Alexa Fluor 594 donkey anti-rabbit secondary antibody (1:1000 in blocking solution) for 12 h at 4 °C. Finally, the sections were washed, dehydrated and coverslipped. Confocal images were acquired with same settings (10× object) using a Nikon C2 or TIE microscope. The cFos+ cells in the hippocampus were quantified in Imaris (Bitplane) and normalized by the mean number of cFos+ cells in dCA3 in the context exposure group in each batch.

### Slice electrophysiology

Mice were anaesthetized by isoflurane (5%) and then deeply anaesthetized by high-dose isoflurane (∼150 µL in a custom nose mask). Mice were transcardially perfused with ice-cold NMDG-based slicing artificial cerebral spinal fluid (ACSF, 93 mM NMDG, 2.5 mM KCl, 1.2 mM NaH2PO4, 30 mM NaHCO3, 20 mM HEPES, 5 mM Sodium ascorbate, 2 mM thiourea, 3 mM Sodium pyruvate, 25 mM D-glucose,12 mM N-Acetyl-L-cysteine, 10 mM MgCl2, 0.5 mM CaCl2, oxygenated with 95% O2/5% CO2). After the perfusion, the brain was immediately removed and transferred to ice-cold NMDG-based slicing ACSF. Coronal brain slices (300 µm thick) containing hippocampus were prepared using a vibratome (VT- 1200S, Leica) in an ice-cold NMDG-based ACSF. Slices were maintained for 12 min at 32°C in NMDG-based ACSF and were subsequently transferred into HEPES-based solution containing (in mM: 92 NaCl, 2.5 KCl,1.2 NaH2PO4, 30 NaHCO3, 20 HEPES, 5

Sodium ascorbate, 2 thiourea, 3 Sodium pyruvate, 25 D-glucose, 2 MgCl2, 2 CaCl2, bubbled with 95% O2/5% CO2) at room temperature and incubated more than 1 h before recording, and then were kept at room temperature (20-22°C) until start of recordings. For the recording, slices were transferred to a recording chamber and infused with ∼30 °C recording ACSF (in mM: 119 NaCl, 5 KCl, 1.25 NaH2PO4, 26 NaHCO3, 10 D-glucose, 1 MgCl2, 2 CaCl2, bubbled with 95% O2/5% CO2). All chemicals were purchased from Sigma-Aldrich. Patch pipettes (4 – 7 MΩ) pulled from borosilicate glass (Sutter instrument, BF150-86-10) were filled with a K-gluconate based internal solution (in mM: 126 K-gluconate, 2 KCl, 2 MgCl2,10 HEPES, 0.2 EGTA, 4 MgATP2, 0.4 Na3GTP, 10 Na-phosphate creatine, 290 mOsm, adjusted to pH 7.2∼7.3 with KOH). Whole-cell recording was performed with a Multiclamp 700B amplifier and a Digidata 1440A (Molecular Device). The Voltage-clamp holding potential and Current-clamp membrane potential were both -70 mV. For optogenetic stimuli, the blue light pulse (480 nm) from a LED engine (Sola SE5-LCR-V8, Lumencor) was triggered by digital commands from the Digidata 1440A and delivered through the objective to illuminate the field of view.

### Comparing the preference for pathway-specific synaptic connections

After the AAV injections into dCA2/3 or iCA3, the ChR2-expressing axons were stimulated by blue light onto vCA1-containing brain slice with TTX (1 µM) bath applied. Patch-clamp recording were made from cells in vCA1d, vCA1m and vCA1v. To compare the synaptic strength of distinct pathways, the optogenetic stimulation setting (∼2 ms, 0.5 mW) was kept constant for the same brain slice.

When measuring synaptic connections of dCA2/3 axons in the same brain slice, at least one cell was recorded from vCA1d and one cell from vCA1m or vCA1v. For each animal, all the light-evoked synaptic currents were normalized by the mean amplitude of all vCA1d neurons recorded.

When measuring synaptic connections of iCA3 axons in the same brain slice, at least one cell was recorded from vCA1m and one cell from vCA1d or vCA1v. For each animal, all the light-evoked synaptic currents were normalized by the mean amplitude of all vCA1m cells recorded.

### Measurements of excitability of vCA1d neurons by dCA2/3 inputs

The LED light (∼5 ms, 0.5 mW) was delivered to the brain slice via the 60x objective lens and hence activated the dCA2/3 axons in vCA1d. For each cell recorded, the light stimuli with increasing intensities were set based on the minimal light intensity that evokes detectable synaptic responses. Each light intensity was repeated for 4 trials and the next intensity was increased in a 0.1 mw step.

### Measurements of excitatory and inhibitory conductances

Voltage-clamp whole-cell recording was performed with varying holding potentials (-70, - 40 & +10 mV) to measure *Isyn* versus *Vm*. The excitatory and inhibitory conductances, *Ge(t)* and *Gi(t)*, were separated by the method adapted from Wehr and Zador(Wehr and Zador, 2003) and computed by

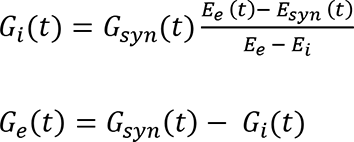

Where the total synaptic conductance *Gsyn(t)* and total reversal potential *Esyn(t)* were obtained from the slope and intercept, respectively, of the linear regression of *Isyn* versus *Vm* (Fig. 5f-h). *Ee* and *Ei* are the reversal potentials for excitatory and inhibitory inputs, respectively. For our Cs-based internal solution (in mM: 125 Cs-gluconate, 8 KCl, 10 HEPES, 1 EGTA, 4 Mg.ATP, 0.5 Na3GTP, 10 Na-phosphate creatine, 5 QX-314, 285∼295 mOsm, 7.2∼7.3 pH), *Ee* = 0 mV and *Ei* = -72.8 mV.

To optogenetically evoke synaptic conductances in vCA1d neurons, the power of LED light was set at minimal intensity that reliably evokes detectable currents. To quantify the *Ge* and *Gi*, a 50 ms window before the stimulation onset was defined as the baseline period and a 1000 ms window after was defined as the response period. The amplitude was computed by the difference between the response peak and baseline mean. The delay was computed by the first time point when the response exceeds one standard deviation (SD) of baseline period. Because of the duration of optogenetic pulse is typically at 2 ms, this conductance delay should be considered as a delay for detectable synaptic conductances instead of a delay for synaptic transmission.

### Contextual fear conditioning (CFC)

Two different contexts were used for CFC experiments. Context A (in cm, 30 [L] × 30 [W] × 35 [H]) has a mental grid floor (Coulbourn Instruments) and clear Plexiglas walls and is cleaned with 70% ethanol. Context B (same size as context A) has a flat acrylic floor and walls with checkboard-pattern papers and is cleaned with 1% acetic acid. The foot shocks were delivered by an animal shocker (H13-15, Coulbourn Instruments). These behavioral contexts were placed in a sound-attenuating chamber. Mice was handled for 5-7 days before CFC. The behavior protocol was controlled by an Anilab system (Anilab, China). The behavior video was recorded by a webcam throughout the paradigm. Freezing behavior was evaluated by a method based on video-frame-pixel changes (scripts available at https://github.com/XuChunLab/FreezingAnalysis). In brief, every 4^th^ – 7^th^ frame in each video was extracted from behavioral videos for freezing evaluation. Pixel values in each frame were converted to gray value (0-255). Pixels that differed more than 30 in consecutive frames were considered as changed pixels. Consecutive frames were considered as immobile episodes if the proportion of change pixels is less than the threshold (constant for the same type of video). The immobile episodes longer than 1 s were marked as freezing episodes and the freezing was expressed as a percentage of time spent freezing. All parameters were optimized based on the behavioral video and kept the same for the same conditioning contexts. These analyzed results were highly correlated with the human quantification and were in the same level of correlation between human experimenters.

*CFC with optogenetics:* For three-trial CFC, 3 foot-shocks were delivered after 3 min exposure to context A (0.7 mA, 2 s, 1 min interval) on day 1. Mice were put back to home cage 1 min after the last shock. On day 2, mice were placed in context A for 5 min to measure the conditioned freezing. On day 3, mice were place in the context B for 5 min to measure the generalized freezing. For one-trial CFC, mice received 1 shock (0.7 mA, 2 s) after 5 min exposure to context A on day 1 and were put back to home cage 1 min after the shock. On day 2, mice were placed into context A for 5 min to measure the conditioned freezing. The same procedures on day 1 and 2 were repeated on day 3 and 4, respectively.

*CFC for slice electrophysiology:* For C57 mice, animal went through three-trial CFC on day 1. Next day, mice were placed in context A for 5 min. For control mice, foot shocks were omitted. For *cFos-tTA* mice, the animal food was replaced by the Doxycycline-containing (DOX, 40 mg/g) food (HY-N0565B, MedChemExpress) right after the AAV injection. Four weeks later, the DOX food was taken off two days before the cell tagging in context A and put on right after the tagging. Mice were randomly assigned to two groups. One week later, one group of mice underwent three-trial CFC in context A and the other group in context B. On the next day, the slice recording was performed on pyramidal cells after contextual fear retrieval in the conditioned context.

*CFC with photometry:* For the behavioral protocol in Fig. 4c, mice were fear conditioned by three foot-shocks (0.7 mA, 2 s, 1 min interval) after 10 min baseline period, and continued to stay in the conditioned context for 30 min post conditioning period. For the behavioral protocol in Fig. 4i, mice were fear conditioned by three foot-shocks (0.7 mA, 2 s, 1 min interval) after 3 min baseline period and continued to stay in the conditioned context for 1 min. The next day, mice were placed back to the conditioned context for 5 min fear retrieval. For the behavioral protocols in Fig. S11, mice were subjected to two days of fear conditioning by 6 foot-shocks (0.7 mA, 2 s, 1 min interval) in order to collect more trials with shock responses. In each animal, there are at least 2-4 channels recorded simultaneously.

*CFC with miniscope:* The day before CFC day, mice were exposed to context A and B for 5 min in each. Some mice from the same batch of mice underwent an additional day of context exposures to A and B in order to do the control analysis without CFC impact. On the CFC day, mice were fear conditioned in context A by three foot-shocks (0.7 mA, 2 s, 1 min interval) after 3 min baseline period. Mice were placed back to home cage 1 min after the last shock. The day after CFC, mice were exposed to context A and B for 5 min, respectively.

### *In vivo* electrophysiology and analysis

*Surgery and single-unit recording:* Mice were anesthetized with isoflurane and head-fixed in a stereotaxic frame (RWD, China). The skull was gently cleaned by 1/10 H2O2 and washed by PBS. Sterile skull screws were drilled into the skull, fixed with Vetbond (3M) and connected with copper wire for reference and grounding. Holes for electrode implantation were drilled above dorsal hippocampus. A custom-built 16-channel tetrode consisting of gold-plated insulated nichrome wires (diameter 10 µm, impedance 0.5 – 1 MΩ; California Fine Wire) was slowly implanted at the coordinates: AP, -1.7, LAT ±2.1, DV -1.7. The end of each wire was deinsulated and attached to a 20-pin connector. The implant was secured on the skull using dental acrylic (Refine Bright, Yamahachi Dental Meg) and protected by copper tape. Mice were allowed to recover for a week after implantation.

The digital headstage (Plexon) was connected with the electrode, and then connected to a 16-channel OmniPlex D-DHP Acquisition System (Plexon) via lightweight data cable. Neuronal activity was digitized at 40 kHz and low-cut Bessel filtered at 250 Hz. One week after the surgery, the hippocampal tetrodes were moved in small daily increments (50 µm) toward CA3 until clear and stable spikes presented. At the conclusion of the experiment, recording sites were marked with electrolytic lesions (20 μA, 20 s) and reconstructed with standard histological procedures.

Single-unit recording is performed when animals explore two contexts (20[L] × 20[W] × 30 cm[H]) with distinct tactile, olfactory and visual information. Context A has square patches on the wall and stainless-steel floor and is cleaned with 75% ethanol. Context B has zebra stripes on the wall and acrylic floor and is cleaned with 1% acetic acid. The door of the context is manually opened and closed after animals freely enter the door. For each trial, the animals explore the context for 5 min before being taken out. In one session, animals typically explore context A and B 8 trials in total (trial interval >1 min) with random order. In separate-context scenario, two contexts are connected with a narrow corridor, from where animals enter into either context. In morph-context scenario, the two contexts have exactly the same location in the room, but the wall, floor and odor of the context are manually changed as needed during the trial interval.

*Spike sorting:* Single-unit spikes were sorted by Offline Spike Sorter (OFSS, Plexon) (Fig. S9). Principal component scores were calculated for unsorted waveforms and plotted on three-dimensional principal component spaces. Discrete clusters with similar waveforms were manually defined and separated by autocorrelation and cross-correlation functions and other criteria including refractory period (1 ms), multivariate ANOVA (p < 0.05), high J3 values (the ratio of between-cluster to within-cluster scatter) and low DB (Davies-Bouldin validity index, the ratio of the sum of within-cluster scatter to between-cluster separation) values.

*Context-modulated cell*: The context modulated units were determined by a bootstrap analysis adapted from Rozeske et al.(Rozeske et al., 2018). For each unit, all the spikes of all trials in two contexts were used to create a surrogate distribution of expected spikes for each context by shuffling the inter-spike intervals from the original spike timestamps (10,000 iterations). Those units that fell outside of the surrogate distribution (P < 0.01) are considered to be context-modulated cell.

*Context discrimination score*: The discrimination score between two different contexts is computed by (RA-RB)/(RA+RB) where RA and RB are mean firing rate across trials in context A and B, respectively. The rate difference index is computed as discrimination score for every cell in each animal. For context A vs. A and B vs. B, 4 trials are randomly divided into two groups to calculate the discrimination score. The rate difference index is computed as the mean of discrimination scores from 3 iterations.

### Photometric recording

Four to six weeks after stereotaxic injections, animals were implanted with optic fibers (NA 0.37, 200 µm Φ, Anilab, China) ∼300 µm above the Ca^2+^ imaging areas (in mm: dCA2/3: AP -1.5, LAT ±2.1, DV -1.4; vCA1d: AP -3.2, LAT ±3.8, DV -1.25, angle 14°; basolateral amygdala: AP -1.2, LAT ±3.0, DV -4.0). Animals were singly housed and were allowed one week for recovery after the surgery. Animals were habituated to the experimenter by handling for at least 5 days beforehand, and the behavioral experiments were conducted during the animal’s light cycle. The optic fibers were connected with a 465 nm blue LED light source (Cinelyzer, Plexon Inc). The excitation light at the tip of fiber was adjusted to be 14-16 µW (setting 10-12 in the LED controller). Emission signals and behavioral videos (640 x 480 pixels at 30 Hz) were captured simultaneously by a multi-fiber photometry recording system (Cinelyzer). When combining with optogenetics, the optical fibers were implanted above the dCA2/3 and connected with a 462 nm diode laser (2 ms pulse, 19-23 mW at the fiber tip, Shanghai lasers, www.lasercentury.com). The trigger of photometry recording and optogenetic pulses were all controlled by Anilab system.

Raw data of emission signals were acquired and directly extracted for analysis in Matlab (MathWorks) without any filtering. For foot shocks, the 2 s before foot shock onset was defined as the baseline and the 10 s period after stimulus onset was defined as the response window. For optogenetic stimulating pulses, the 1 s before light pulse onset was defined as the baseline and the 2 s period after stimulus onset was defined as the response window. The baseline mean was computed as F0. After removing the artifact of light stimulation pulses (2-5 ms) in dCA2/3, the Ca^2+^ signals in the response window were normalized by the baseline ([F-F0]/F0). The area under the curve (a.u.c.) of Ca^2+^ signals in the response window were then quantified for the event-related responses.

### Optogenetics in behaving animals

For optogenetic manipulations of behavior, optic fibers (200 µm Φ, NA 0.37, Anilab, China) were bilaterally implanted in the vCA1 (vCA1d: AP -3.2mm, LAT 3.4mm, DV -1.2mm; vCA1m: AP -3.2mm, LAT 3.4mm, DV -3.3mm) four weeks after stereotaxic injections. Optic fibers were fixed on the skull with light curing dental resin (CharmFil Flow, DentKist Inc) and dental cement (153018AH, Lang Dental Inc, USA). Animals were singly housed and were allowed one week for recovery after the surgery. Animals were handled for 5-7 days before behavioral task. The behavioral experiments were conducted during the animal’s light cycle.

During behavior sessions with optogenetic inhibition, a DPSS yellow laser was connected with optic fibers and was controlled by TTL signals (YL589T6, 589 nm wavelength, Shanghai Lasers, China). The laser intensity was 19 – 23 mW at end of the optic fiber. For optogenetic manipulation during contextual fear conditioning, the light was switched ON 2 s before each shock onset and switched OFF 1 s after the shock offset. For novel context recognition, the light was ON throughout the first exposure (4 min) to a new context (30 cm triangle shape, acrylic floor, cleaned with sodium hypochlorite disinfectant) on day 1. On day 2, animals were placed to the same context again (4 min). The behavioral recording trigger and optogenetic stimuli were controlled by Anilab system. The behavioral tracking was analyzed by the Ethovision XT (Noldus) and further analyzed in by scripts written in Python.

### Miniscope Ca^2+^ imaging

Four to six weeks after AAV injection, a gradient index lens (64519, 1.8 mm Φ, 0.25 pitch, 0.55NA, Edmunt) was implanted above the injection site during a second surgery. Briefly, a small craniotomy was made above the hippocampus and the brain tissue above the target was aspirated with 100 µL pipette attached to a vacuum pump (BT100-1F, LongerPump). PBS was repeatedly applied to the exposed tissue to prevent drying. Aspiration was stopped once a thin layer of horizontal fibers on the hippocampus surface was visualized. Once the surface of hippocampus was clear of blood, the GRIN lens was slowly lowered above the dCA2/3 or vCA1d with a custom holder (RWD, China) in the stereotaxic arm (depth from brain surface, dCA2/3, 1.5 mm; vCA1d, 1.3 mm) and fixed to the skull using light curing dental resin (CharmFil Flow, DentKist Inc). Finally, dental acrylic (Super Bond C&B, Jakarta) was used to seal the skull. One to two weeks after GRIN lens implantation, we started to check for GCaMP6f fluorescence using a miniature epifluorescence microscope (miniscope, UCLA V2, LabMaker). The miniscope was connected to a portable computer for live view of the fluorescence imaging to guide the alignment and focal planes. Once clear field of view with sufficient expression was observed, we fixed the miniscope via a baseplate to the skull using light curing dental resin and dentate acrylic under isoflurane anesthesia. The miniscope was detached, the baseplate sealed with a baseplate cover and the animal returned to its home cage for recovery. Imaging experiments were performed at 5-6 weeks after AAV injections. Mice were habituated to the miniscope mounting procedure for three days before the start of behavioral sessions. The miniscope was mounted on a daily basis right before the behavioral session. The behavioral video recording (Cinelyzer, Plexon Inc.) and the miniscope imaging were synchronized by a Master-8 (A.M.P.I., Israel). Digital imaging data was acquired from CMOS imaging sensor (Aptina, MT9V032), transmitted to the computer via the DAQ box and USB. The Miniscope data was acquired at 30 frames per second and recorded to uncompressed AVI files by DAQ software (MiniScopeControl, UCLA).

### Ca^2+^ imaging analysis

The miniscope data across sessions were concatenated together. The field of view was cropped and then 2x spatially and 3x temporally down-sampled using *moviepy* python package to reduce the computation load afterwards. The preprocessed data was processed by CaImAn (1.8.5) toolbox (Giovannucci et al., 2019) using constrained nonnegative matrix factorization extension (CNMFe) (Zhou et al., 2018). The motion correction and source extraction were all done with default parameter setting except the following: down- sampling factor in time for initialization (tsub) 4; neuron diameter (gSiz) 13 pixels; 2D Gaussian kernel smoothing (gSig) 3 pixels; minimum peak to noise ratio (min_pnr) 10; spatial consistency (rval_thr) 0.85. The accepted raw trace (estimates.C) of neuronal activity was manually inspected in case of abnormal baseline shift. The Ca^2+^ transients were extracted for further analyses using *findpeaks* function (Matlab) with minimum peak prominence larger than two SD of the analysed session. To quantify and compare Ca^2+^ signals across various conditions, the area under the curves for dF/F (%) were computed and normalized by the time duration.

To analyze the shock responses of each cell, the 4 s before shock onset is defined as the baseline period. The shock period (2 s) and the 4 s post shock are together defined as the shock responsive window. The shock responsive (SR) cells were identified by comparing the shock-evoked fluorescence Ca^2+^ signals of individual cells to their baseline fluorescence levels, if the shock-evoked responses were significantly different from baseline activity (Wilcoxon rank-sum test, P < 0.05) and the difference between response mean and baseline mean is larger than one SD of baseline.

To identify the context-modulated cells in miniscope recordings, the spike activity (estimates.S) was deconvolved from raw trace (estimates.C) of Ca^2+^ activity by CNMFe and the context-modulated cells were determined in the same way (bootstrap) as those in single-unit recordings. One animal for CA3 recording was not included due to the lack of deconvolved spike activity after the CNMFe processing.

*Context discrimination score:* To compare neuronal activities for context discrimination between context A (FA) and context B (FB), we compute the FA and FB using the non-freezing episodes during context exposures; because the Ca^2+^ signals were significantly silenced during the freezing episodes (Fig. S12). Those cells with detected Ca^2+^ events in at least one context were included in this analysis. We then compute the raw discrimination score for each cell by (FA-FB)/(FA+FB) where FA and FB are mean Ca^2+^ signals across trials in context A and B, respectively. The absolute value of the raw discrimination score was considered as absolute discrimination score.

*Population vector correlation*: For each animal, the population vector (PoV) was computed from all cells by stacking their mean Ca^2+^ signals (non-freezing episodes) during the context exposure on top of each other (rows) (Danielson et al., 2016; Hainmueller and Bartos, 2018). The PoV correlation between context A and B was then determined as the cross-correlations of these cellular activities, and then was compared before and after CFC.

### Quantification and statistical analysis

The summary of quantification was reported as the mean value with the standard error of the mean (S.E.M.). For the box plot, data are presented as median (center line) with 25/75 percentile (box) and min and max of the data points (whiskers); the plus or filled circle indicates mean. The cell number is indicated with *n* and the animal number is indicated with *N*. No samples were measured repeatedly for statistics. No statistical methods were used to predetermine sample sizes. The sample sizes were chosen based on published studies and current standards in the field.

Statistical analysis was performed in GraphPad Prism 8 or MATLAB. The The D’Agostino & Pearson normality test was tested for all dataset. If the null hypothesis of normal distribution was rejected, non-parametric statistical tests were used. Mann-Whitney U test, Wilcoxon matched-paired signed rank test, two-sided unpaired *t*-test, paired *t*-test, and one-way and two-way ANOVA test were used to test for statistical significance. Statistical parameters including the exact value of n, precision measures (mean ± SEM) and statistical significance are reported in the text or in the figure legends (see individual sections). The significance threshold was *a* = 0.05 (n.s., P > 0.05; *, P < 0.05; **, P < 0.01; ***, P < 0.001; ****, P < 0.0001).

---

**Supplementary Figure 1.**
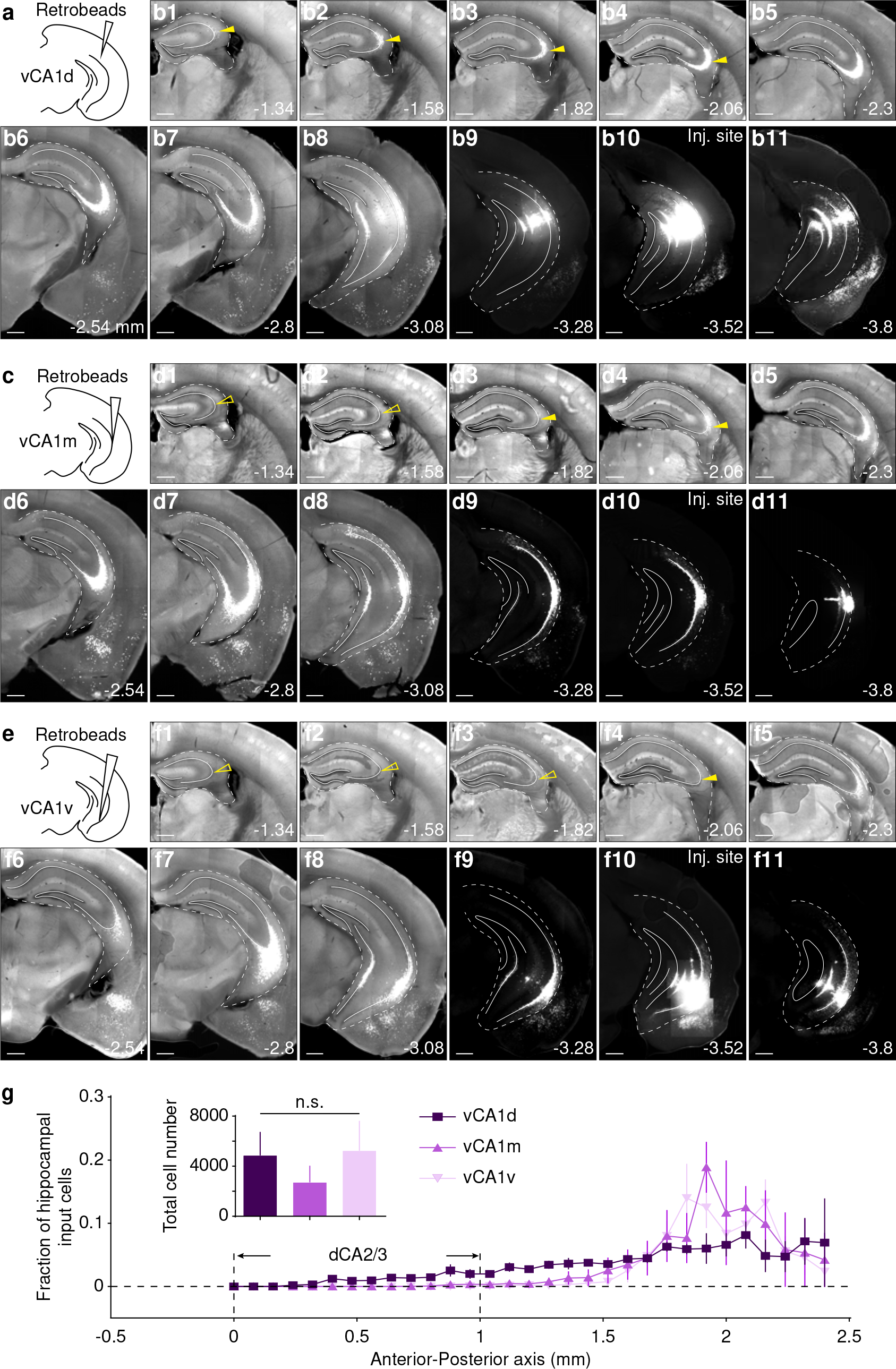
The retrobeads-based retrograde tracing from vCA1 subregions. (a, c, e) Scheme illustrating injection of red retrobeads into vCA1d, vCA1m and vCA1v, respectively. (b, d, f) Examples showing fluorescence of red retrobeads labeled cells across coronal sections of hippocampus. Filled arrows indicated positive labeling by retrobeads and open arrows indicated negative. (g) Summary of proportion of retrobeads labeled input cells in the hippocampus. The anterior-posterior axis is offset by the first dorsal hippocampal section at Bregma -0.94 mm. Inset: summary of total number of retrobeads labeled input cells by retrograde tracing from vCA1d, vCA1m and vCA1v (vCA1d, 4903 ± 1794, N = 3 mice; vCA1m, 2756 ± 1238, N = 3 mice; vCA1v, 5286 ± 2294, N = 4 mice; One-way ANOVA, F(2,7) = 0.46, P = 0.65). Data is summarized as mean ± SEM. Scale bars, 500 µm.

**Supplementary Figure 2.**
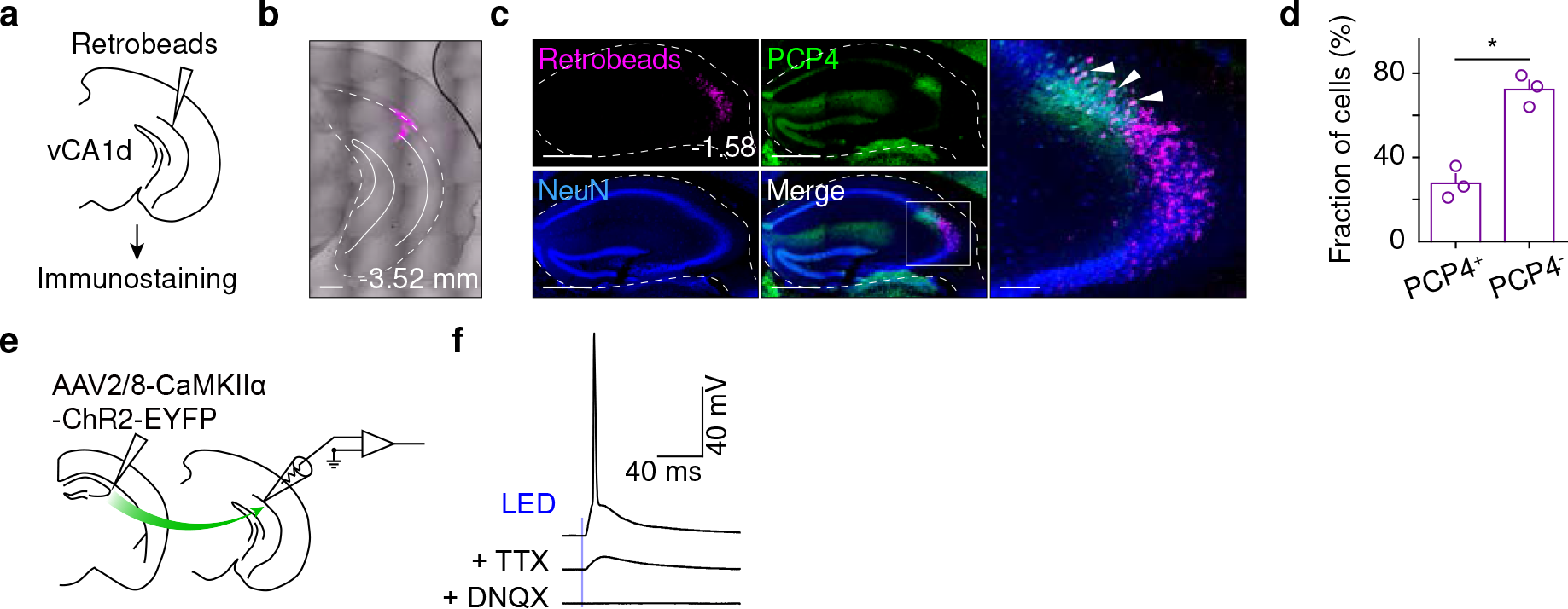
The identity of vCA1d-projecting cells. (a – b) Scheme illustrating injection of red retrobeads into vCA1d and immunostaining example showing red retrobeads at injection site overlaid with image by transmission light. Scale bar, 500 µm. (c) Example showing fluorescence of red retrobeads, immunostaining of PCP4 and NeuN and overlay images (left). Right, the magnification of boxed area on the left. Scale bars, 500 and 100 µm. (d) Percentage summary (mean ± SEM) of PCP4 positive and negative cells in retrobeads labeled cells (27.7 ± 4.4% positive vs. 72.3 ± 4.4% negative; paired *t*-test, P = 0.037, N = 3 mice). (e) Scheme illustrating injection of AAV-CaMKIIα-ChR2 into dCA2/3 and slice recording in vCA1d. (f) Example showing whole-cell recordings of a vCA1d cell upon optogenetic stimulation (blue line) of dCA2/3 axons without and with TTX (1 µM) and DNQX (20 µM) added into the bath sequentially.

**Supplementary Figure 3.**
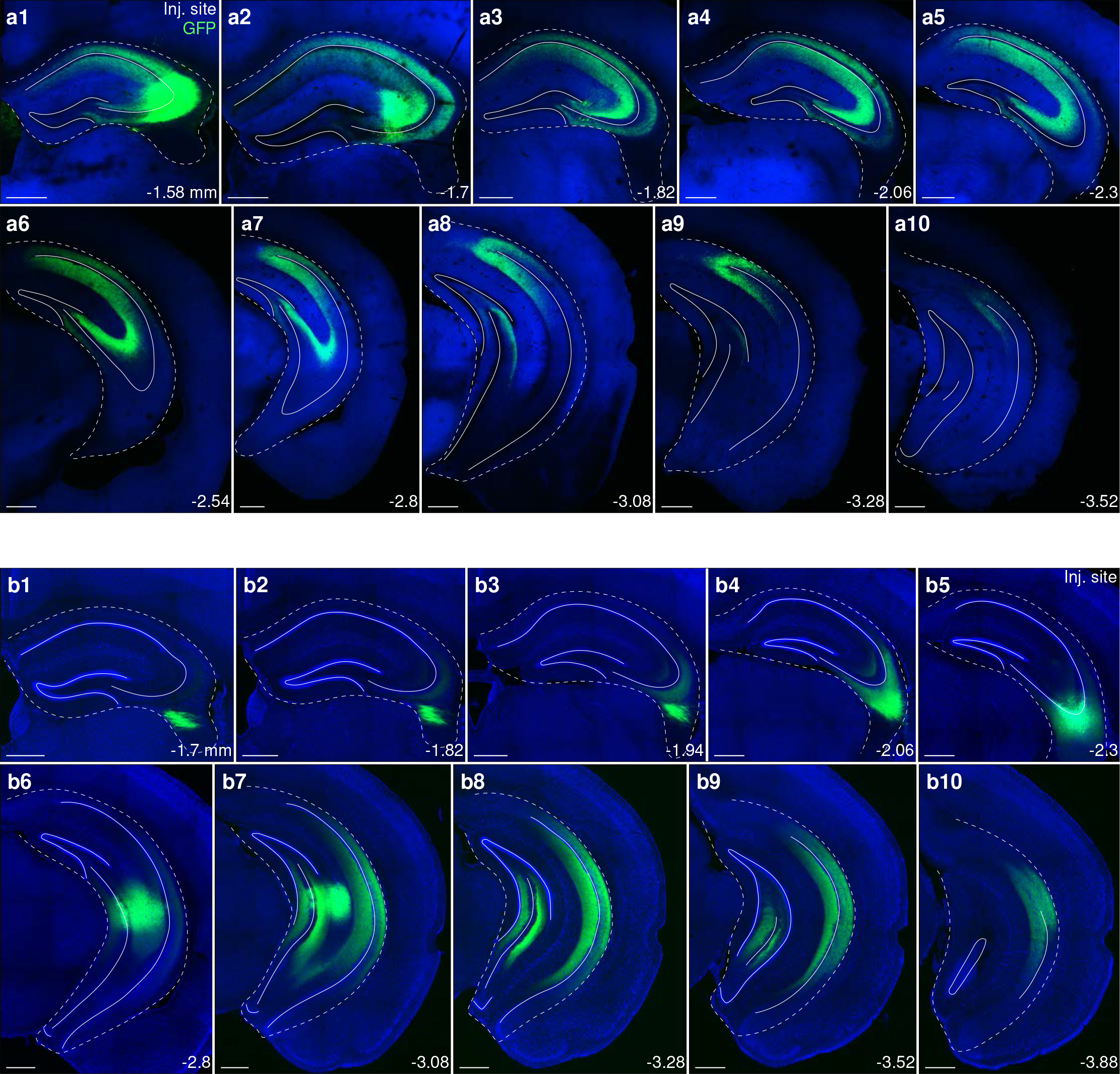
Axon tracing from dCA2/3 neurons versus from iCA3 neurons. (a) Example showing fluorescence of axons from dCA2/3. Injection site at bregma -1.58 mm. (b) Example showing fluorescence of axons from iCA3. Injection site at bregma -2.3 mm. Scale bar, 500 µm.

**Supplementary Figure 4.**
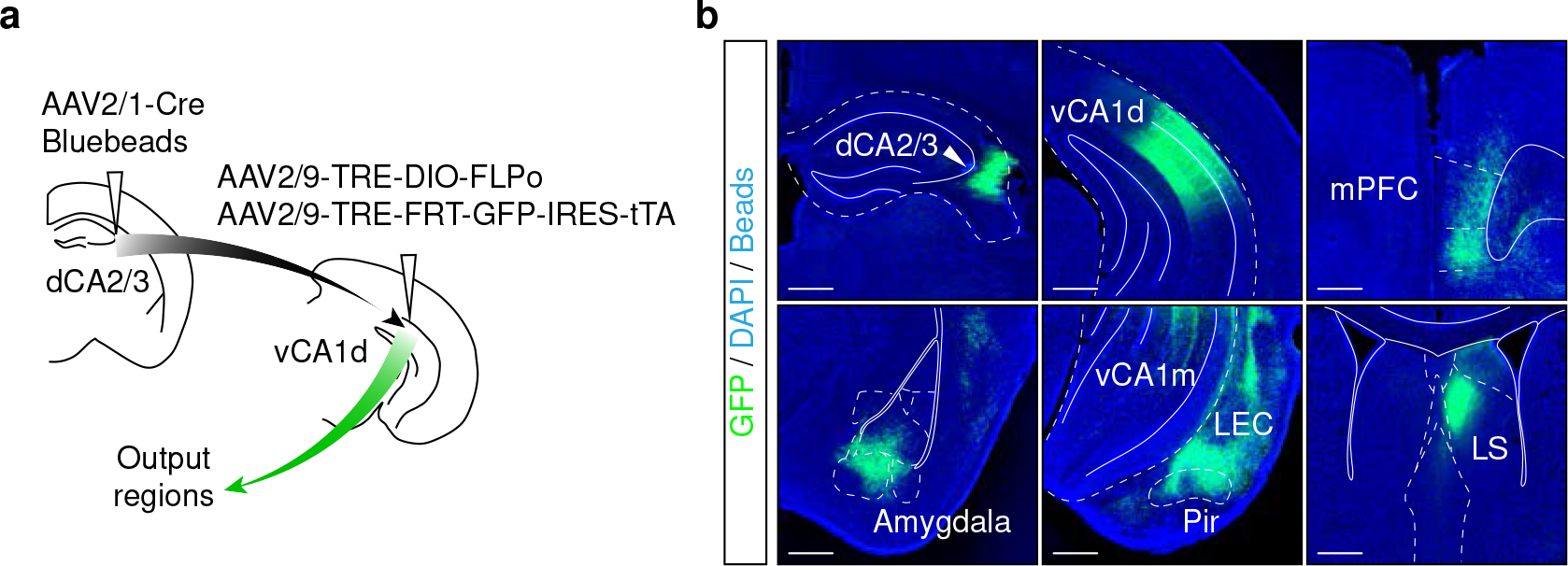
Axon tracing from vCA1d neurons postsynaptic to dCA2/3. (a) ) Scheme for AAV injections in dCA2/3 and vCA1d. (b) Examples showing GFP-expressing axons of vCA1d neurons that received direct inputs from dCA2/3. The injection site in dCA2/3 was marked by blue beads. Scale bar, 500 µm.

**Supplementary Figure 5.**
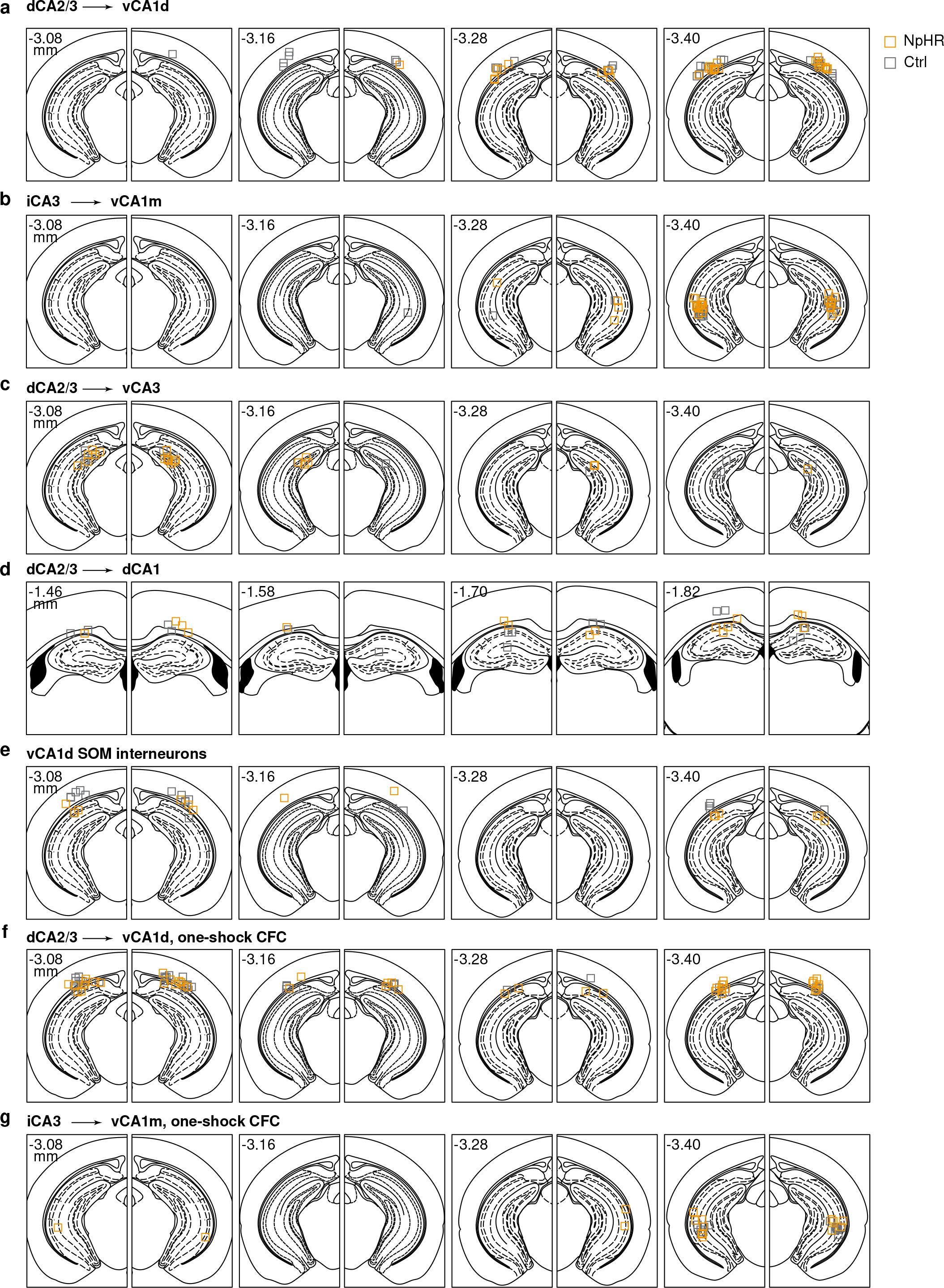
Optical fiber placements for behavior experiments. (a – g) Positions of implanted optical fiber tips (NpHR, yellow; Control, grey) in animals with optogenetic manipulations.

**Supplementary Figure 6.**
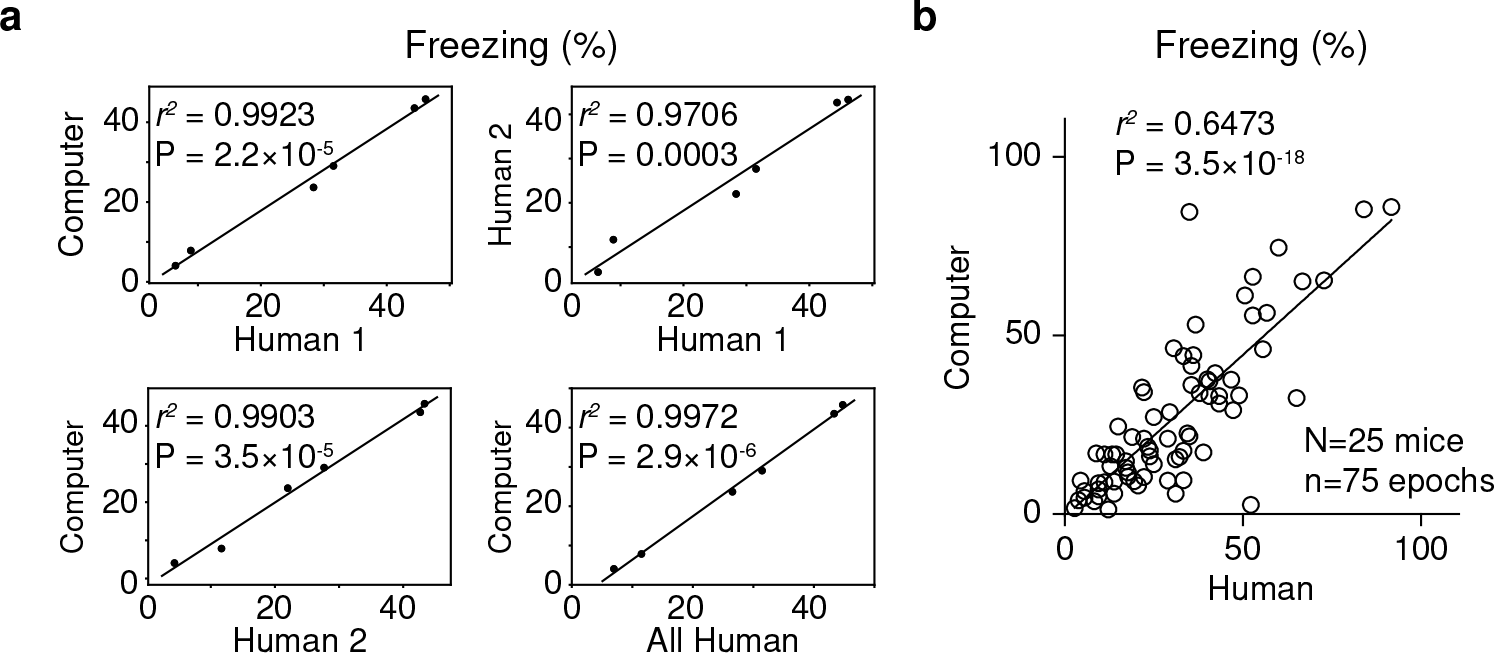
The validation of frame-pixel-based freezing quantification. (a) The correlation (linear regression) among freezing scores (% time spent freezing) for individual animals that were quantified by frame-pixel-based algorithm and human observers. (b) The correlation (linear regression) between the freezing scores for individual epochs (75 freezing epochs from 25 mice) that were quantified by frame-pixel-based algorithm and human observers.

**Supplementary Figure 7.**
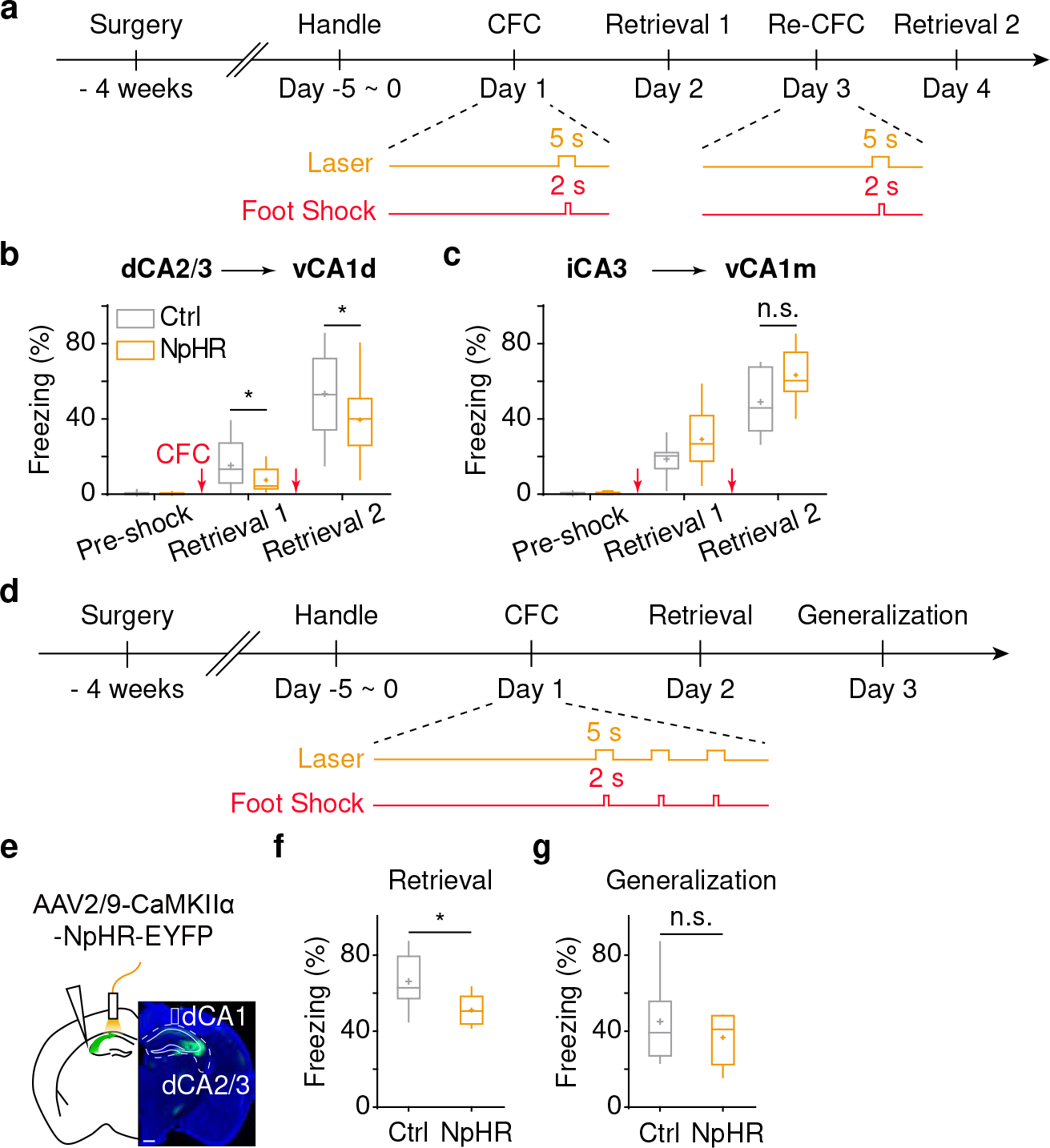
Optogenetic manipulation of hippocampal circuits during CFC. (a) ) Scheme for experimental procedures and behavioral protocols for one-shock CFC. (b) Optogenetic experiments on dCA2/3→vCA1d pathway. The summary (mean ± SEM) of freezing in conditioned context after one-shock CFC (Retrieval 1: 15.3 ± 2.7% Ctrl vs. 7.5 ± 1.3% NpHR; P = 0.012, unpaired *t*-test; Retrieval 2: 53.4 ± 4.6% Ctrl vs. 39.5 ± 4.2% NpHR; P = 0.033, unpaired *t*-test; N = 19 mice for Ctrl, N = 21 mice for NpHR). (c) Optogenetic experiments on iCA3→vCA1m pathway. The summary (mean ± SEM) of freezing in conditioned context after one-shock CFC (Retrieval 1: 18.7 ± 3.1% Ctrl vs. 29.2 ± 6.0% NpHR; P = 0.14, unpaired *t*-test; Retrieval 2: 49.0 ± 5.9% Ctrl vs. 63.3 ± 5.1% NpHR; P = 0.089, unpaired *t*-test; N = 8 mice for Ctrl, N = 8 mice for NpHR). (d) Scheme illustrating experimental procedures. (e) Scheme showing the AAV injection, fiber implantation and example picture of NpHR- EYFP fluorescence in dCA2/3 and dCA1. Scale bar, 500 µm. (f and g) Box plot of freezing in fear retrieval (f) and fear generalization (g) after CFC (Conditioned freezing, 66.2 ± 4.9% Ctrl vs. 51.1 ± 2.8% NpHR; unpaired *t*-test, P = 0.019; Generalized freezing, 45.0 ± 7.4% Ctrl vs. 36.7 ± 4.7% NpHR; unpaired *t*-test, P = 0.36; N = 8 mice for Ctrl, N = 8 mice for NpHR).

**Supplementary Figure 8.**
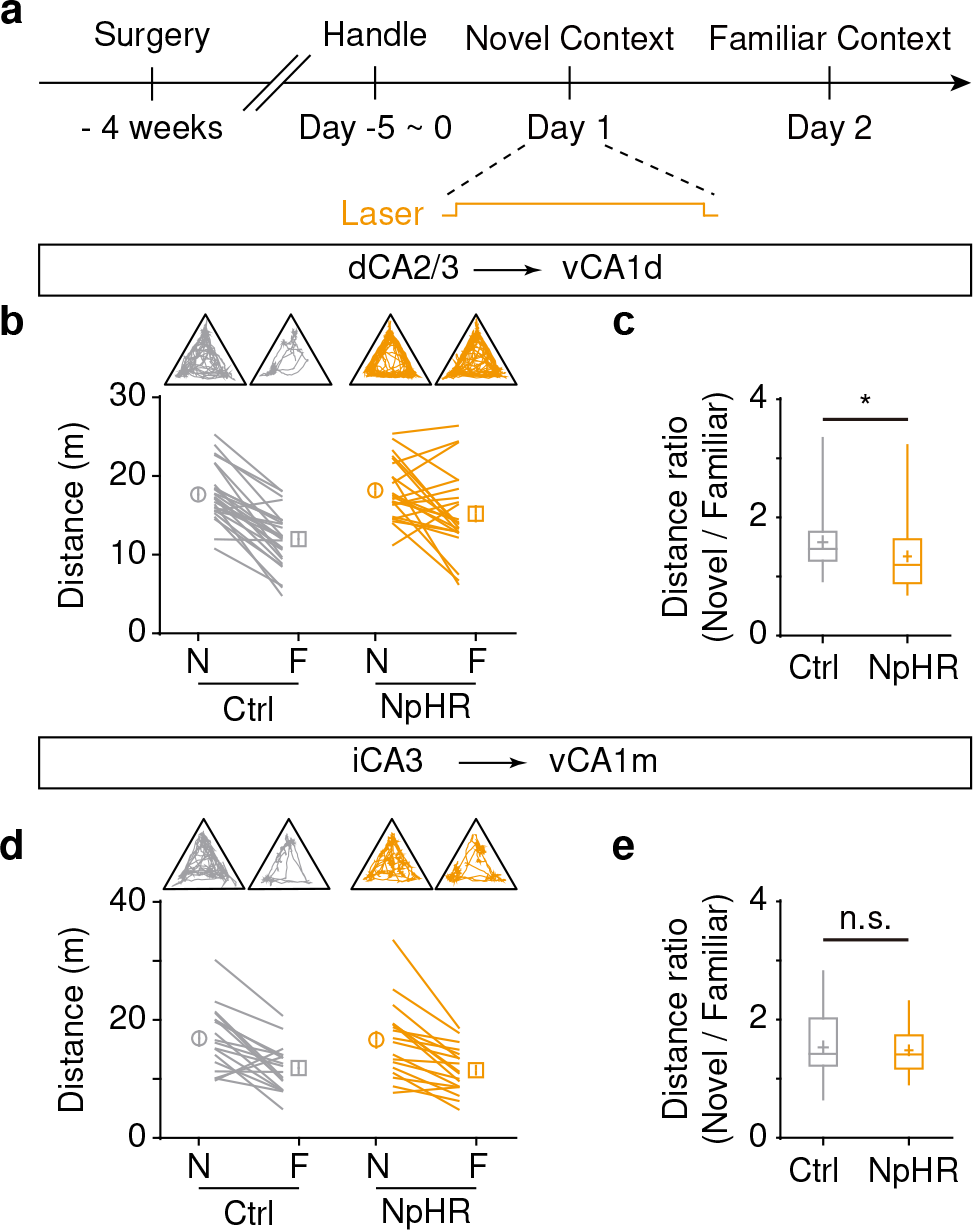
The dCA2/3**→**vCA1d pathway contributes to novel context recognition. (a) Scheme for experimental procedures and behavioral protocols of novel context recognition with optogenetics. (b and c) Optogenetic experiments on dCA2/3→vCA1d pathway. (b) Top, examples showing the behavioral tracking during context exposures on day 1 and day 2 of two animals injected with NpHR (yellow) and Ctrl (grey) AAVs, respectively. Bottom, the summary of travelled distances in two groups of animals. Each line represents one animal. (c) The summary of distance ratio in Ctrl and NpHR groups (Ctrl, 1.6 ± 0.1, N = 27 mice; NpHR, 1.3 ± 0.1, N = 23 mice; Mann-Whitney U test, P = 0.021). (d and e) Optogenetic experiments on iCA3→vCA1m pathway. (d) Examples showing behavioral tracking and the summary of travelled distances. (e) The summary of distance ratio in Ctrl and NpHR groups (Ctrl, 1.5 ± 0.1, N = 18 mice; NpHR, 1.5 ± 0.1, N = 18 mice; Mann-Whitney U test, P = 0.83).

**Supplementary Figure 9.**
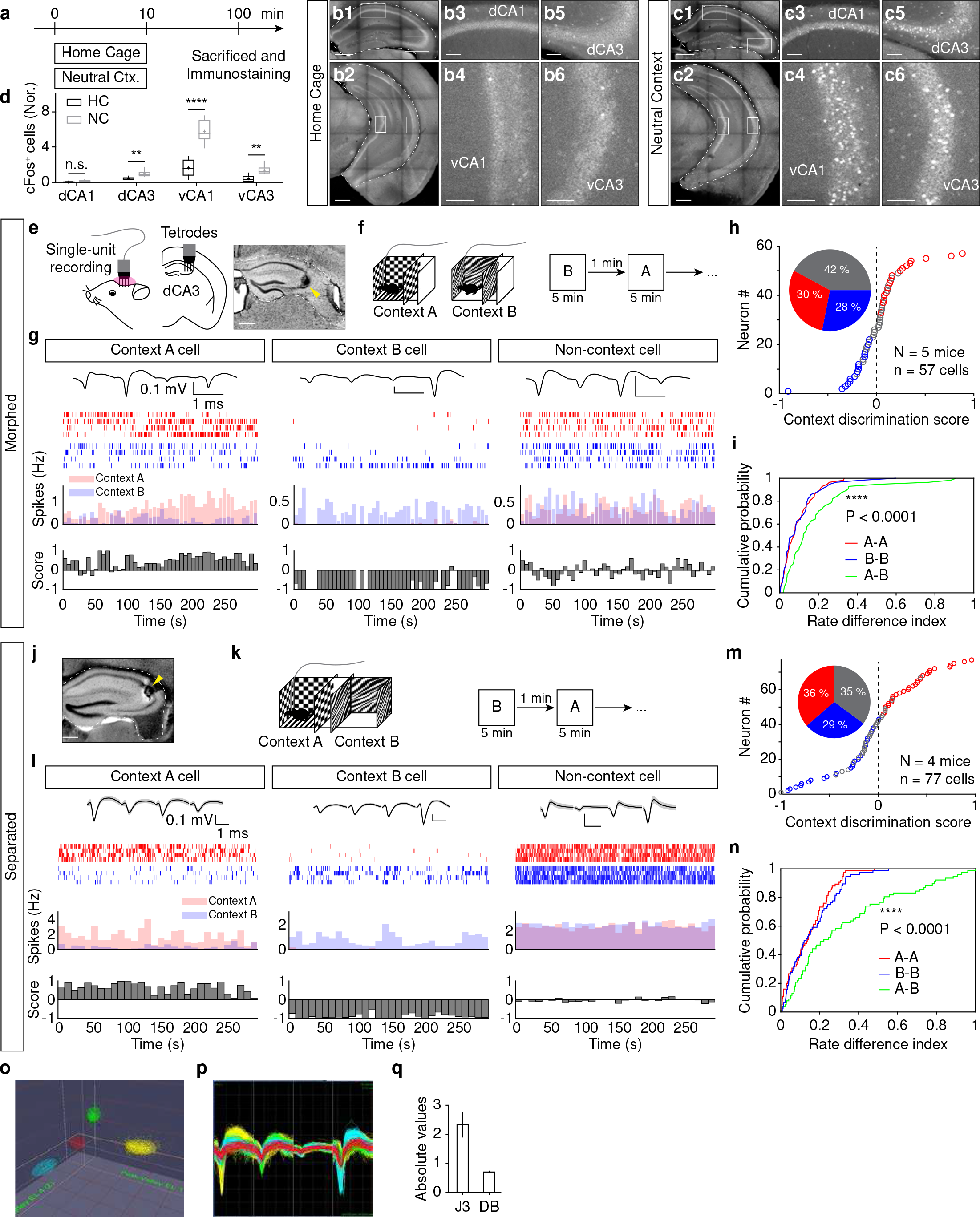
The context-modulated activity in distal part of dCA3. (a) Scheme for experimental procedures for cFos staining. (b and c) Examples showing retrobeads labeling in hippocampal subregions of home cage group and context exposure group. Right, magnification of boxes in left. (d) Summary (mean ± SEM) of cFos+ cells in hippocampal subregions in two groups (normalized by context-activated cFos+ cells in dCA3, grey, 1.0 ± 0.2, N = 6 mice; home cage, black, 0.44 ± 0.1, N = 6 mice; Mann-Whitney U test, P = 0.0043). (e) Left, scheme for tetrodes implantation and single-unit recording in dCA3. Right, histology example showing recording site marked by electrolytic lesion (arrow). (f) Scheme illustrating the behavioral protocol (morphed) for single-unit recording in behaving mice. (g) Example recording of single units identified as context A, context B and non-context preferring cells. Top, example waveforms recorded in the tetrodes. Middle, raster plots of spikes during context exposure in context A (red) and B (blue). One row represents one trial. Bottom, summary (bin = 5 s) of spike rates and computed discrimination scores. (h) Cumulative distribution of context discrimination scores (n = 57 cells from 5 mice). Inset pie chart shows the percentages of context A (17/57), context B (16/57) and non- context preferring cells (24/57). (i) Cumulative probability of the rate difference indexes computed from mean spike rates in distinct contexts (A-B, shown in green) versus same contexts (A-A, shown in red; B-B, shown in blue). Kolmogorov-Smirnov (KS) test: A-A vs. A-B, P = 9.60 × 10^-14^; B-B vs. A-B, P = 9.60 × 10^-14^; A-A vs. B-B, P = 0.98. (j) Histology example showing recording site marked by electric lesion (arrow). (k) Scheme illustrating the behavioral protocol (separated) with two-connected separate contexts for single-unit recording in behaving mice. (l) Example recording of single units identified as context A, context B and non-context preferring cells. Top, example waveforms recorded in the tetrodes. Middle, raster plots of spikes during context exposure in context A (red) and B (blue). Bottom, summary (bin = 5 s) of spike rates and computed discrimination scores. (m) Cumulative distribution of context discrimination scores (n = 77 cells from 4 mice). Inset shows the pie chart of percentages of context A (28/77), context B (22/77) and non- context preferring cells (27/77). (n) Cumulative probability of the rate difference indexes computed from the mean spike rate in distinct contexts versus same contexts. Kolmogorov-Smirnov test, A-A vs. A-B, P = 5.61×10^-5^, B-B vs. A-B, P = 4.49×10^-4^, A -A vs. B-B, P = 0.65. (o) Spikes from individual units were sorted into cluster by 3D principal component analysis. (p) Superimposed waveforms of four units recorded from the same tetrode in dCA2/3. (q) Recording quality was accessed by J3 and Davies Bouldin validity index (DB) statistics. High J3 and low DB values indicate good isolation of multiple units. Scale bars, 500 and 100 µm.

**Supplementary Figure 10.**
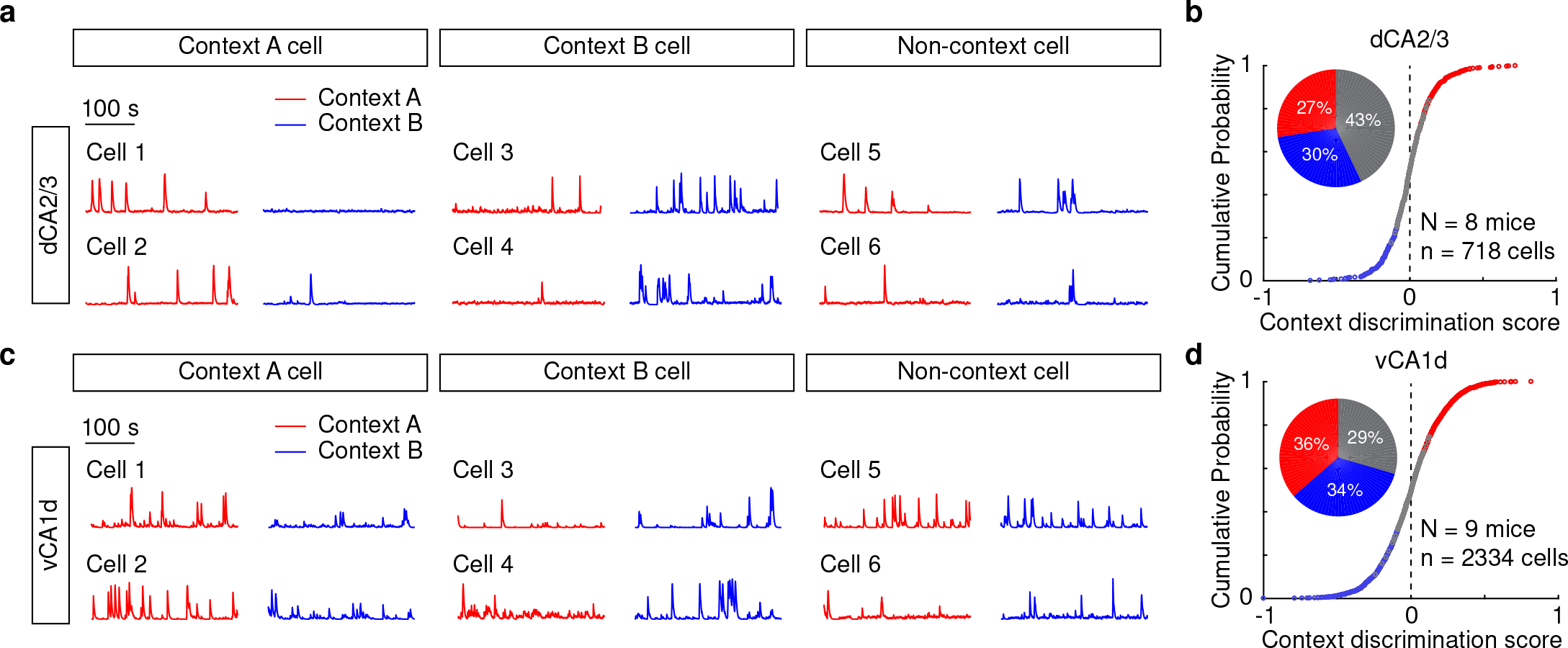
Context-modulated Ca^2+^ activity of dCA2/3 and vCA1d neurons. (a and c) Example showing Ca^2+^ activity of individual neurons in the dCA2/3 (a) and vCA1d (c) identified as context A, context B and non-context preferring cells. (b and d) Cumulative distribution of context discrimination scores. Inset pie chart shows the percentages of context A, context B and non-context preferring cells in the dCA2/3 (b) and vCA1d (d).

**Supplementary Figure 11.**
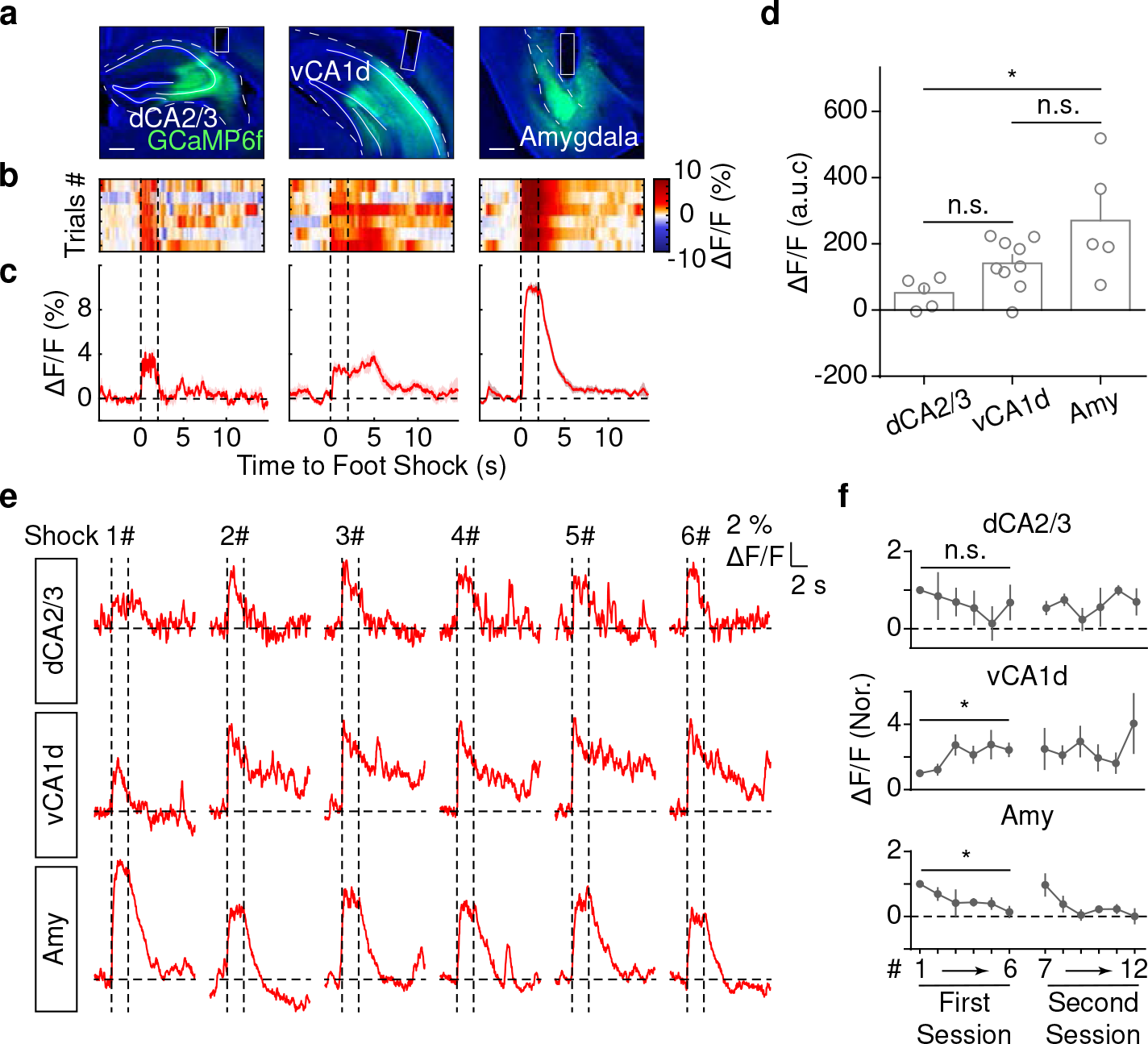
The photometric Ca^2+^ responses to foot shocks in dCA2/3, vCA1d and amygdala. (a) Example showing GCaMP6f fluorescence and fiber tracks in dCA2/3, vCA1d and amygdala. (b) Heatmap of Ca^2+^ signals upon foot shock (between dash lines, 2s, 0.7 mA). (c) ) Mean traces of 6 trials of foot shocks. (d) Shock-evoked response (a.u.c.) in dCA2/3, vCA1d and amygdala. (dCA2/3, 52.3 ± 20.4; vCA1d, 141.4 ± 25.5; amygdala, 270.9 ± 77.6; One-way ANOVA, F(2,16) = 5.541, P = 0.015; Turkey’s multiple comparisons test, dCA2/3 vs. amygdala, P = 0.012; dCA2/3, N = 5 mice; vCA1d, N = 9 mice; amygdala, N = 5 mice). (e) Example of photometric Ca^2+^ signals upon foot shocks (vertical lines, 0.7 mA, 2 s, 1 min interval). (f) Summary (mean ± SEM, normalized by the response to 1^st^ shock) of Ca^2+^ signals in dCA2/3, vCA1d and amygdala when animal went through 2 sessions of foot shocks in two days (Paired *t*-test; dCA2/3: 0.68 ± 0.44, N = 3 mice; P = 0.88; vCA1d: 2.43 ± 0.39, N = 4 mice; P = 0.046; amygdala: 0.14 ± 0.16, N = 4 mice; P = 0.011). Scale bars, 500 µm.

**Supplementary Figure 12.**
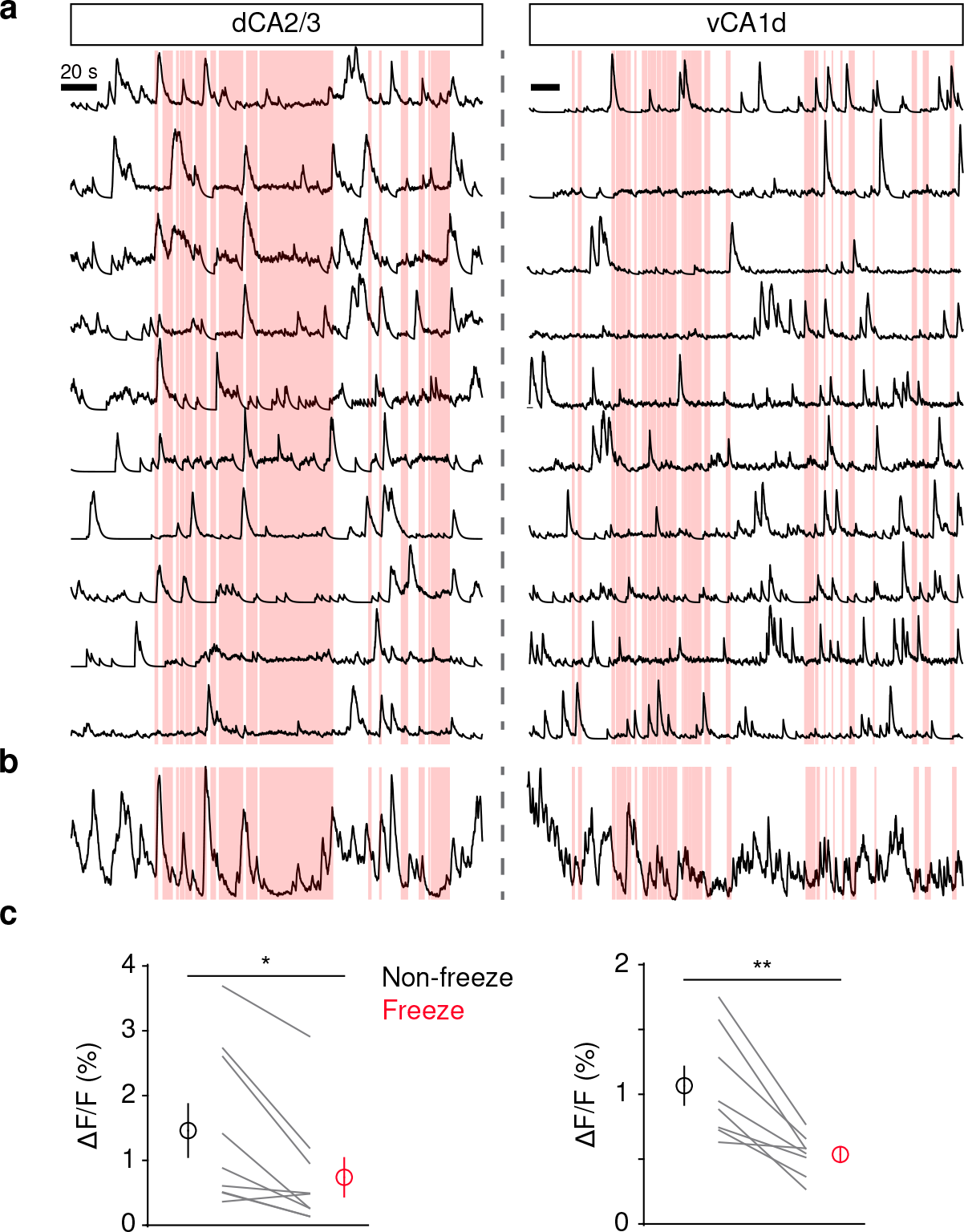
The suppression of Ca^2+^ activity in dCA2/3 and vCA1d during freezing episodes. (a) Example traces showing Ca^2+^ signals during the fear retrieval in the conditioned context (Left, one animal with dCA2/3 imaging; right, one animal with vCA1d imaging). The red shadings indicate the freezing episodes. (b) The averaged traces from all cells imaged from the two animals in (a) (n = 82 cells for dCA2/3; n = 503 cells for vCA1d). (cC) Summary (mean ± SEM) of Ca^2+^ signals during freezing vs. non-freezing episodes in dCA2/3 and vCA1d. Each line represents one animal. 1.48 ± 0.41 % Non-freezing vs. 0.76 ± 0.30 % Freezing; Wilcoxon signed rank test, P = 0.012; N = 9 mice for dCA2/3. 1.07 ± 0.15 % Non-freezing vs. 0.54 ± 0.056 % Freezing; paired *t*-test, P = 0.003; N = 8 mice for vCA1d.

**Supplementary Figure 13.**
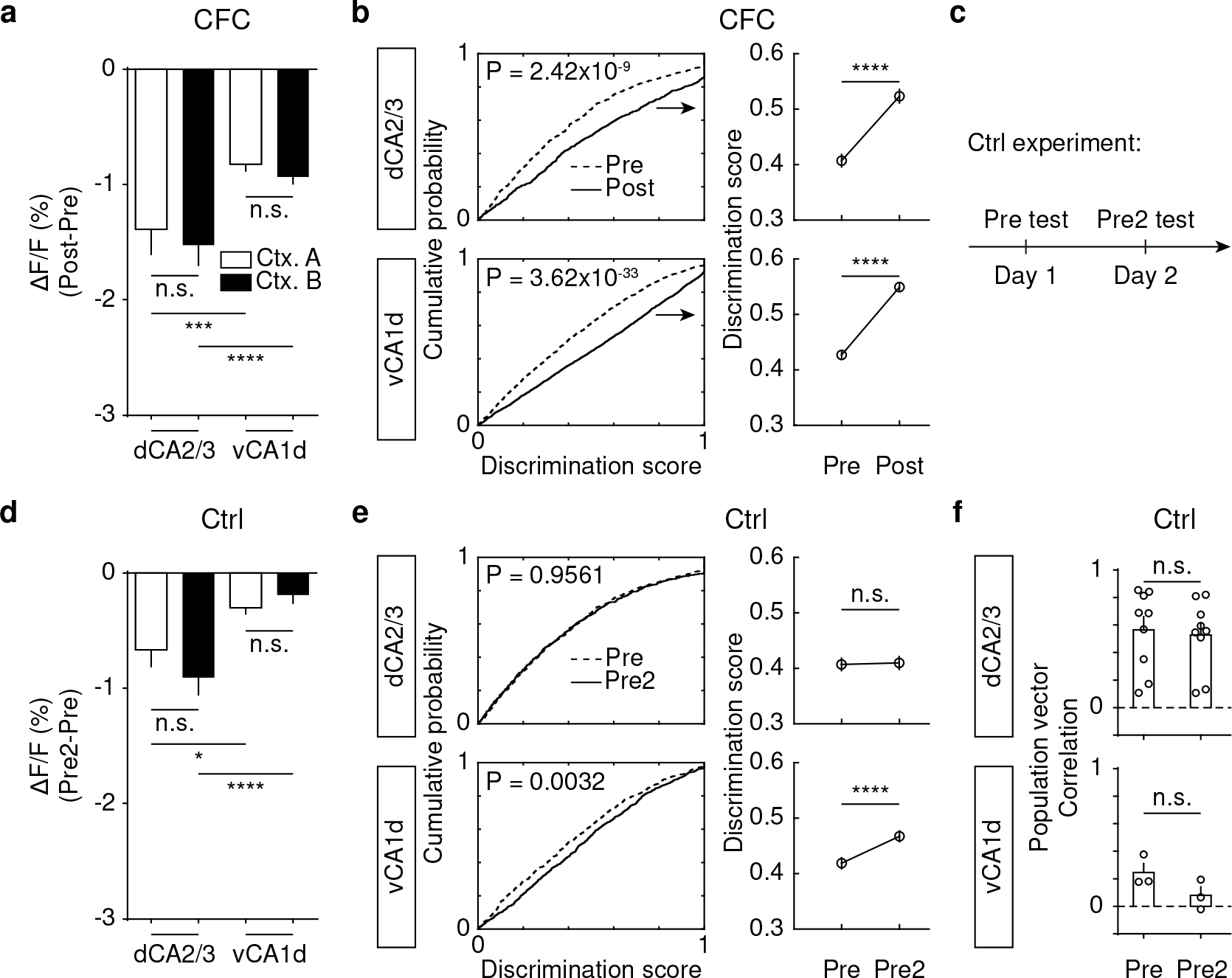
The context-evoked Ca^2+^ activity in dCA2/3 and vCA1d from animals with and without CFC. (a) Summary of post-CFC changes in context-evoked Ca^2+^ signals of dCA2/3 and vCA1d neurons (dCA2/3 vs. vCA1d, unpaired *t*-test: Ctx. A, -1.39 ± 0.22% vs. -0.82 ± 0.07%, P = 0.0009; Ctx. B, -1.53 ± 0.18% vs. -0.93 ± 0.06%, P = 4.44×10^-5^). (b) Left, the cumulative distribution of absolute context discrimination scores of dCA2/3 and vCA1d neurons before and after CFC (Kolmogorov-Smirnov test, dCA2/3, P = 2.42×10^-9^; vCA1d, P = 3.62×10^-33^). Right, the summary of discrimination scores (Pre vs. Post, paired *t*-test: dCA2/3, 0.41 ± 0.01 vs. 0.52 ± 0.01, P = 1.12×10^-13^; vCA1d, 0.43 ± 0.01 vs. 0.55 ± 0.01, P = 3.51×10^-54^). (c) ) Scheme for the behavioral protocol without shocks. (d) Summary of changes in context-evoked Ca^2+^ signals of dCA2/3 and vCA1d neurons during the first (Pre) and second (Pre2) context exposure without shocks (dCA2/3 vs. vCA1d, unpaired *t*-test: Ctx. A, -0.67 ± 0.15% vs. -0.30 ± 0.053%, P = 0.019; Ctx. B, -0.91 ± 0.15% vs. -0.19 ± 0.072%, P = 1.50×10^-5^). (e) Left, the cumulative distribution of absolute context discrimination scores of dCA2/3 and vCA1d neurons during the first (Pre) and second (Pre2) context exposure without foot shocks (Kolmogorov-Smirnov test, dCA2/3, P = 0.9561; vCA1d, P = 0.0032). Right, the summary of discrimination scores (Pre vs. Pre2, paired *t*-test: dCA2/3, 0.41 ± 0.01 vs. 0.41 ± 0.01, P = 0.85; vCA1d, 0.42 ± 0.01 vs. 0.47 ± 0.01, P = 5.40×10^-6^). (f) Summary of population vector correlation of neuronal activity between context A and B in mice without CFC (Pre vs. Pre2, paired *t*-test; dCA2/3, 0.56 ± 0.1 vs. 0.53 ± 0.09, N = 9 mice, P = 0.59; vCA1d, 0.25 ± 0.07 vs. 0.08 ± 0.06, N = 3 mice, P = 0.30). Data is summarized as mean ± SEM.

**Supplementary Figure 14.**
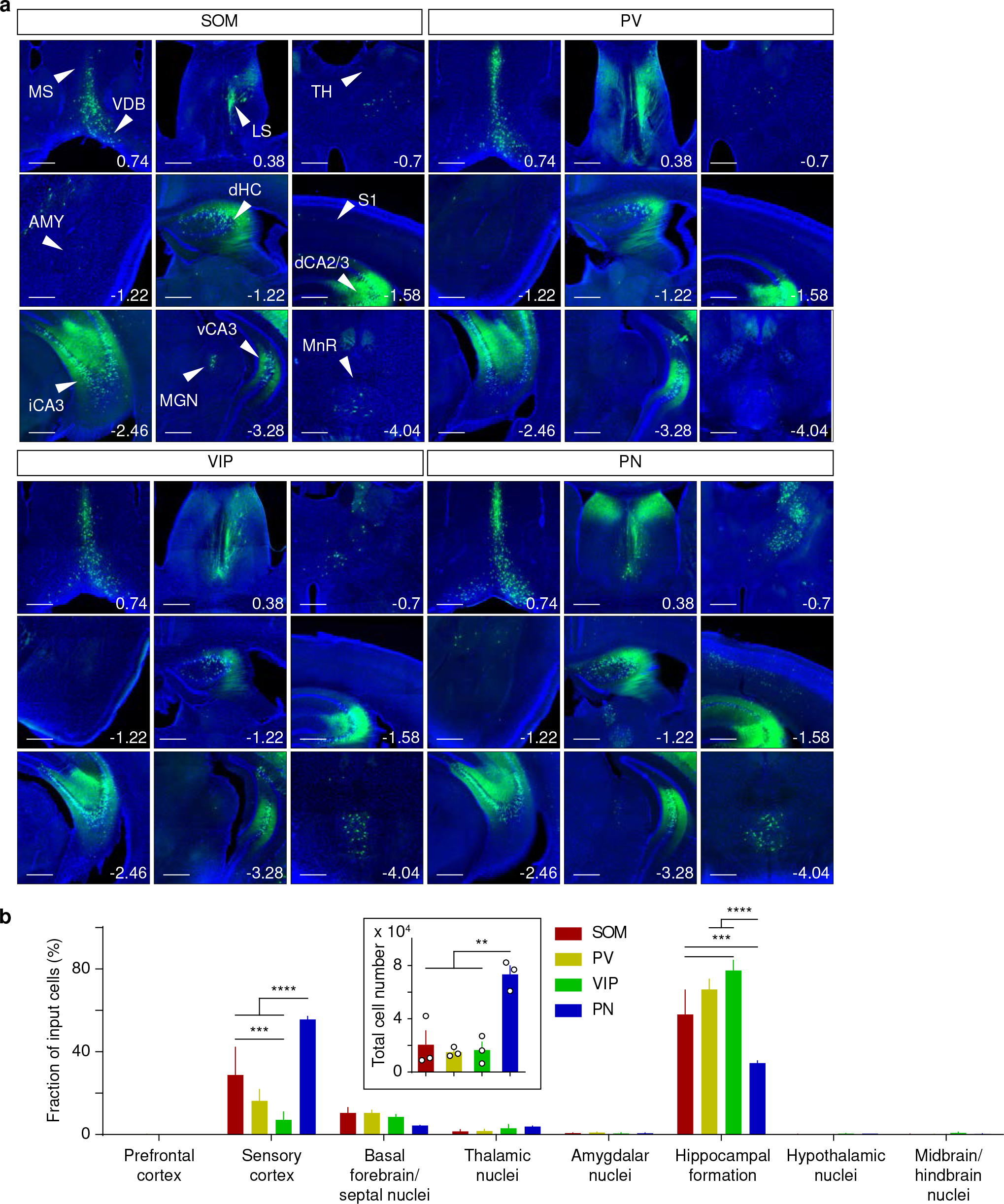
Cell-type specific tracing from subtypes of vCA1d neurons. (a) Examples showing rabies-labeled presynaptic cells to specific cell types in vCA1d. Scale bars, 500 µm. (b) Summary (mean ± SEM) of proportion of brain-wide presynaptic cells in upstream brain areas. Inputs from sensory cortex, SOM: 28.8 ± 13.4%, N = 3 mice; PV: 16.3 ± 5.5%, N = 3 mice; VIP: 7.1 ± 3.8%, N = 3 mice; PN: 55.6 ± 1.5%, N = 3 mice; Two-way ANOVA, F(21,64) = 8.764, P = 7.0×10^-12^; Turkey’s multiple comparisons test, PN vs. SOM, P = 1.6×10^- 5^; PN vs. PV, P = 1.2×10^-9^; PN vs. VIP, P = 1.1×10^-11^; SOM vs. VIP, P = 5.3×10^-4^; Inputs from hippocampus, SOM: 58.0 ± 11.8%; PV: 70.1 ± 5.0%; VIP: 79.3 ± 4.9%; PN: 34.5 ± 1.1%; Turkey’s multiple comparisons test, PN vs. SOM, P = 1.6×10^-4^; PN vs. PV, P = 2.1×10^-8^; PN vs. VIP, P = 2.5×10^-11^; SOM vs. VIP, P = 6.8×10^-4^. Inset, total number of presynaptic cells. SOM: 20511 ± 10830, N = 3 mice; PV: 14950 ± 2009, N = 3 mice; VIP: 16564 ± 5968, N = 3 mice; PN: 73387 ± 6400, N = 3 mice; One-way ANOVA, F(3,8) = 15.98, P = 0.001; Turkey’s multiple comparisons test, PN vs. SOM, P = 0.0032; PN vs. PV, P = 0.0017; PN vs. VIP, P = 0.002.

## References

1. Amaral, D.G., and Witter, M.P. (1989). The three-dimensional organization of the hippocampal formation: a review of anatomical data. Neuroscience 31, 571–591.

2. Andersen, P., Bliss, T.V., and Skrede, K.K. (1971). Lamellar organization of hippocampal pathways. Exp Brain Res 13, 222–238.

3. Bast, T., Zhang, W.N., and Feldon, J. (2001). The ventral hippocampus and fear conditioning in rats. Different anterograde amnesias of fear after tetrodotoxin inactivation and infusion of the GABA(A) agonist muscimol. Exp Brain Res 139, 39–52.

4. Cai, D.J., Aharoni, D., Shuman, T., Shobe, J., Biane, J., Song, W., Wei, B., Veshkini, M., La-Vu, M., Lou, J., et al. (2016). A shared neural ensemble links distinct contextual memories encoded close in time. Nature 534, 115–118.

5. Danielson, N.B., Zaremba, J.D., Kaifosh, P., Bowler, J., Ladow, M., and Losonczy, A. (2016). Sublayer-Specific Coding Dynamics during Spatial Navigation and Learning in Hippocampal Area CA1. Neuron 91, 652–665.

6. Fanselow, M.S. (2000). Contextual fear, gestalt memories, and the hippocampus. Behav Brain Res 110, 73–81.

7. Fanselow, M.S., and Dong, H.W. (2010). Are the dorsal and ventral hippocampus functionally distinct structures? Neuron 65, 7–19.

8. Fanselow, M.S., and Poulos, A.M. (2005). The neuroscience of mammalian associative learning. Annual review of psychology 56, 207–234.

9. Gabernet, L., Jadhav, S.P., Feldman, D.E., Carandini, M., and Scanziani, M. (2005). Somatosensory integration controlled by dynamic thalamocortical feed-forward inhibition. Neuron 48, 315–327.

10. Giovannucci, A., Friedrich, J., Gunn, P., Kalfon, J., Brown, B.L., Koay, S.A., Taxidis, J., Najafi, F., Gauthier, J.L., Zhou, P., et al. (2019). CaImAn an open source tool for scalable calcium imaging data analysis. Elife 8.

11. Goshen, I., Brodsky, M., Prakash, R., Wallace, J., Gradinaru, V., Ramakrishnan, C., and Deisseroth, K. (2011). Dynamics of retrieval strategies for remote memories. Cell 147, 678–689.

12. Gradinaru, V., Zhang, F., Ramakrishnan, C., Mattis, J., Prakash, R., Diester, I., Goshen, I., Thompson, K.R., and Deisseroth, K. (2010). Molecular and cellular approaches for diversifying and extending optogenetics. Cell 141, 154–165.

13. Hainmueller, T., and Bartos, M. (2018). Parallel emergence of stable and dynamic memory engrams in the hippocampus. Nature 558, 292–296.

14. Hippenmeyer, S., Vrieseling, E., Sigrist, M., Portmann, T., Laengle, C., Ladle, D.R., and Arber, S. (2005). A developmental switch in the response of DRG neurons to ETS transcription factor signaling. PLoS Biol 3, e159.

15. Hitti, F.L., and Siegelbaum, S.A. (2014). The hippocampal CA2 region is essential for social memory. Nature 508, 88–92.

16. Jimenez, J.C., Berry, J.E., Lim, S.C., Ong, S.K., Kheirbek, M.A., and Hen, R. (2020). Contextual fear memory retrieval by correlated ensembles of ventral CA1 neurons. Nat Commun 11, 3492.

17. Jung, M.W., Wiener, S.I., and McNaughton, B.L. (1994). Comparison of spatial firing characteristics of units in dorsal and ventral hippocampus of the rat. J Neurosci 14, 7347–7356.

18. Kheirbek, M.A., Drew, L.J., Burghardt, N.S., Costantini, D.O., Tannenholz, L., Ahmari, S.E., Zeng, H., Fenton, A.A., and Hen, R. (2013). Differential control of learning and anxiety along the dorsoventral axis of the dentate gyrus. Neuron 77, 955–968.

19. Kim, J.J., and Fanselow, M.S. (1992). Modality-specific retrograde amnesia of fear. Science 256, 675–677.

20. Kim, W.B., and Cho, J.H. (2020). Encoding of contextual fear memory in hippocampal-amygdala circuit. Nat Commun 11, 1382.

21. Kjelstrup, K.B., Solstad, T., Brun, V.H., Hafting, T., Leutgeb, S., Witter, M.P., Moser, E.I., and Moser, M.B. (2008). Finite scale of spatial representation in the hippocampus. Science 321, 140–143.

22. LeDoux, J.E. (2000). Emotion circuits in the brain. Annu Rev Neurosci 23, 155–184.

23. Lee, H., Wang, C., Deshmukh, S.S., and Knierim, J.J. (2015). Neural Population Evidence of Functional Heterogeneity along the CA3 Transverse Axis: Pattern Completion versus Pattern Separation. Neuron 87, 1093–1105.

24. Leutgeb, J.K., Leutgeb, S., Moser, M.B., and Moser, E.I. (2007). Pattern separation in the dentate gyrus and CA3 of the hippocampus. Science 315, 961–966.

25. Lin, J.Y., Lin, M.Z., Steinbach, P., and Tsien, R.Y. (2009). Characterization of engineered channelrhodopsin variants with improved properties and kinetics. Biophys J 96, 1803–1814.

26. Liu, X., Ramirez, S., Pang, P.T., Puryear, C.B., Govindarajan, A., Deisseroth, K., and Tonegawa, S. (2012). Optogenetic stimulation of a hippocampal engram activates fear memory recall. Nature 484, 381–385.

27. Lovett-Barron, M., Kaifosh, P., Kheirbek, M.A., Danielson, N., Zaremba, J.D., Reardon, T.R., Turi, G.F., Hen, R., Zemelman, B.V., and Losonczy, A. (2014). Dendritic inhibition in the hippocampus supports fear learning. Science 343, 857–863.

28. Lu, L., Igarashi, K.M., Witter, M.P., Moser, E.I., and Moser, M.B. (2015). Topography of Place Maps along the CA3-to-CA2 Axis of the Hippocampus. Neuron 87, 1078–1092.

29. Maren, S. (2001). Neurobiology of Pavlovian fear conditioning. Annu Rev Neurosci 24, 897–931.

30. Maren, S., Phan, K.L., and Liberzon, I. (2013). The contextual brain: implications for fear conditioning, extinction and psychopathology. Nat Rev Neurosci 14, 417–428.

31. Nakazawa, K., Quirk, M.C., Chitwood, R.A., Watanabe, M., Yeckel, M.F., Sun, L.D., Kato, A., Carr, C.A., Johnston, D., Wilson, M.A., and Tonegawa, S. (2002). Requirement for hippocampal CA3 NMDA receptors in associative memory recall. Science 297, 211–218.

32. O’Keefe, J., and Nadel, L. (1978). The Hippocampus as a Cognitive Map (Oxford University Press).

33. Phillips, R.G., and LeDoux, J.E. (1992). Differential contribution of amygdala and hippocampus to cued and contextual fear conditioning. Behavioral neuroscience 106, 274–285.

34. Pitkanen, A., Pikkarainen, M., Nurminen, N., and Ylinen, A. (2000). Reciprocal connections between the amygdala and the hippocampal formation, perirhinal cortex, and postrhinal cortex in rat. A review. Annals of the New York Academy of Sciences 911, 369–391.

35. Reijmers, L.G., Perkins, B.L., Matsuo, N., and Mayford, M. (2007). Localization of a stable neural correlate of associative memory. Science 317, 1230–1233.

36. Rozeske, R.R., Jercog, D., Karalis, N., Chaudun, F., Khoder, S., Girard, D., Winke, N., and Herry, C. (2018). Prefrontal-Periaqueductal Gray-Projecting Neurons Mediate Context Fear Discrimination. Neuron 97, 898–910 e896.

37. Scoville, W.B., and Milner, B. (1957). Loss of recent memory after bilateral hippocampal lesions. J Neurol Neurosurg Psychiatry 20, 11–21.

38. Strange, B.A., Witter, M.P., Lein, E.S., and Moser, E.I. (2014). Functional organization of the hippocampal longitudinal axis. Nat Rev Neurosci 15, 655–669.

39. Swanson, L.W., and Cowan, W.M. (1977). An autoradiographic study of the organization of the efferent connections of the hippocampal formation in the rat. J Comp Neurol 172, 49–84.

40. Swanson, L.W., Wyss, J.M., and Cowan, W.M. (1978). An autoradiographic study of the organization of intrahippocampal association pathways in the rat. J Comp Neurol 181, 681–715.

41. Taniguchi, H., He, M., Wu, P., Kim, S., Paik, R., Sugino, K., Kvitsiani, D., Fu, Y., Lu, J., Lin, Y., et al. (2011). A resource of Cre driver lines for genetic targeting of GABAergic neurons in cerebral cortex. Neuron 71, 995–1013.

42. Tovote, P., Fadok, J.P., and Luthi, A. (2015). Neuronal circuits for fear and anxiety. Nat Rev Neurosci 16, 317–331.

43. Wagatsuma, A., Okuyama, T., Sun, C., Smith, L.M., Abe, K., and Tonegawa, S. (2018). Locus coeruleus input to hippocampal CA3 drives single-trial learning of a novel context. Proc Natl Acad Sci U S A 115, E310–E316.

44. Wehr, M., and Zador, A.M. (2003). Balanced inhibition underlies tuning and sharpens spike timing in auditory cortex. Nature 426, 442–446.

45. Wickersham, I.R., Lyon, D.C., Barnard, R.J., Mori, T., Finke, S., Conzelmann, K.K., Young, J.A., and Callaway, E.M. (2007). Monosynaptic restriction of transsynaptic tracing from single, genetically targeted neurons. Neuron 53, 639–647.

46. Xu, C., Krabbe, S., Grundemann, J., Botta, P., Fadok, J.P., Osakada, F., Saur, D., Grewe, B.F., Schnitzer, M.J., Callaway, E.M., and Luthi, A. (2016). Distinct Hippocampal Pathways Mediate Dissociable Roles of Context in Memory Retrieval. Cell 167, 961–972 e916.

47. Yang, S., Yang, S., Moreira, T., Hoffman, G., Carlson, G.C., Bender, K.J., Alger, B.E., and Tang, C.M. (2014). Interlamellar CA1 network in the hippocampus. Proc Natl Acad Sci U S A 111, 12919–12924.

48. Zhang, F., Wang, L.P., Brauner, M., Liewald, J.F., Kay, K., Watzke, N., Wood, P.G., Bamberg, E., Nagel, G., Gottschalk, A., and Deisseroth, K. (2007). Multimodal fast optical interrogation of neural circuitry. Nature 446, 633–639.

49. Zhou, H., Xiong, G.J., Jing, L., Song, N.N., Pu, D.L., Tang, X., He, X.B., Xu, F.Q., Huang, J.F., Li, L.J., et al. (2017). The interhemispheric CA1 circuit governs rapid generalisation but not fear memory. Nat Commun 8, 2190.

50. Zhou, P., Resendez, S.L., Rodriguez-Romaguera, J., Jimenez, J.C., Neufeld, S.Q., Giovannucci, A., Friedrich, J., Pnevmatikakis, E.A., Stuber, G.D., Hen, R., et al. (2018). Efficient and accurate extraction of in vivo calcium signals from microendoscopic video data. Elife 7.

